# Evolution and plasticity of the transcriptome under temperature fluctuations in the fungal plant pathogen *Zymoseptoria tritici*

**DOI:** 10.1101/725010

**Authors:** Arthur J. Jallet, Arnaud Le Rouzic, Anne Genissel

## Abstract

**– Background.** Species are subjected to variable environment in nature. While the ability to cope with changes has been studied through theoretical approaches, the small number of empirical studies limits our understanding of the evolutionary molecular mechanisms underlying adaptation to fluctuations **– Results.** Whole transcriptome sequencing of the fungal wheat pathogen *Zymoseptria tritici* revealed the evolution of gene expression after 48 weeks of experimental evolution under stable or fluctuating temperature. We found that although there is a strong genetic signature of gene expression, fluctuating regime could have a large and pervasive effect on the transcriptome evolution. Results show a few hot spots of significantly differentially expressed genes in the genome, in regions enriched with transposable elements (TE), including on dispensable chromosomes. Further, our results evidenced gene evolution towards robustness associated with the fluctuating regime, suggesting robustness is adaptive in changing environment. Last, an analysis on gene expression correlation revealed a significant set of genes that could act as trans-regulators. **– Conclusions.** This study is the first evolve and re-sequence experiment of a fungal pathogen, with the goal to describe the potential ability to evolve to changing environment and the molecular changes occurring in response to fluctuating selection. In addition to explore the evolutionary potential of the pathogen under temperature fluctuation this work highlights the important role of transcriptional rewiring that can adjust regulation of cell growth and multiplication.

## Background

One central theme in evolutionary genetics is to understand the role of regulatory variants in the molecular basis of adaptation. In particular in the context of rapid global climate change with growing evidence that species experience more abiotic fluctuations, it becomes more important to study the role of regulatory variants in adaptation to changing environments. Since early arguments in favor of regulatory variation explaining phenotypic differences between organisms (Britten and Davidson, 1971; King and Wilson, 1975), over the past two decades or so empirical studies have found abundant variation of gene expression within and between species (*e.g.* in Primates (Enard et al., 2002); in teleost fish (Oleksiak et al., 2002; Whitehead and Crawford, 2006); in Humans (Rockman and Wray, 2002); in Drosophila (Whittkop et al., 2004; and Saccharomyces (Fay et al., 2004); and evidenced the widespread of heritable variation of gene expression within and between populations, much of which is thought to be adaptive (Brem and Kruglyak, 2002; Leder et al, 2015; Noormohammad et al., 2017; Glaser-Schmitt and Parsh, 2018).

Extensive body of work has shown that environmental change causes rapid gene expression response (*e.g. Caenorhabditis elegans* (Li et al., 2006; Feugeas et al., 2016). With regard to temperature, many studies have evidenced large and pervasive effect on the transcriptome of the temperature, paying much attention on cold and hot stresses (*i.e.*, Chen et al., 2015; Jovic et al., 2017; Zhou et al., 2012), including temperature fluctuations (Podrasbky and Somero, 2004; Sørensen et al., 2016). Testing the gene expression variation in variable environments also raises the question of the role of the phenotypic plasticity in adaptive evolution. Plasticity is the ability of a genotype to produce different phenotypes in different environments (Pigliuici et al., 2006). Evidence of adaptive plasticity that contributes first to the establishment of adaptation remains a debate (see Ghalambor et al., 2007; Levis and Pfenning, 2016) though mechanisms of adaptation by mean of plasticity were nicely demonstrated in a few recent landmark papers (Corl et al., 2018; Ghalambor et al., 2015; Kenkel and Matz, 2016).

One powerful approach to understand how gene expression evolves in the face of environmental variation is experimental evolution, which consists of laboratory-controlled evolution that lasts over several generations (Garland and Rose, 2009). Premises to examine the molecular responses to environmental fluctuations has been fulfilled using experimental evolution in *Drosophila melanogaster* using diet fluctuations (Huang and Agrawal, 2016; Zanveld et al., 2017); in *Escherichia coli* long term evolution under changing temperature (Bennett, Lenski and Mittler 1992; Leroi, Lenski and Bennett 1994), variable pH (Hughes et al., 2007) and variable abiotic stresses in *Saccharomyces cerevisiae* (Dhar et al., 2013). In contrast, there are too few studies on filamentous fungi (Fisher and Lang, 2016).

In the present study, we investigate the transcriptome evolution of the wheat fungal pathogen *Zymoseptoria tritici*, to assess the effect of temperature fluctuations on gene expression evolution and gene expression plasticity. Like other micro-organisms, *Z. tritici* displays clear advantages for experimental evolution, due to its short generation time, small genome and the ease of maintenance in the laboratory. *Z. tritici* is a filamentous Ascomycete fungus, and is the main causal agent of the Septoria Tritici Blotch (STB) disease on wheat (O’Driscoll et al., 2014; Fones and Gurr, 2015). *Z. tritici* is a haploid single-cell species that multiply asexually *in vitro* by budding (Steinberg, 2015). Nowadays this species becomes a good fungal model with growing interest to study its genome evolution since the publication of its complete reference genome (Goodwin et al., 2011). While much attention has been paid towards the ability of the pathogen to overcome the immune system of its host through QTL mapping approach (Meile et al., 2018; Stewart et al., 2018) or Genome Wide Association Studies (Zhong et al., 2017; Hartmann et al., 2017), we still have a poor understanding of the potential ability of *Z. tritici* to adapt to abiotic changes (Zhan and McDonald, 2011; Lendemann et al., 2016), and we have very poor knowledge on the contribution of regulatory variants to adaptation for this fungal species.

Transcriptome studies in the fungal pathogen *Z. tritici* are fairly recent and aimed for the most part at characterizing the waves of up- and down-regulated genes in association with symptom development inside the host plant (Kellner et al., 2014; Rudd et al., 2015; Palma-Guerrero et al., 2017). The present study is the first analysis of transcriptome evolution in response to temperature fluctuations in this pathogenic fungus. We used an evolve-and-resequence approach to trace the changes of gene expression that occurred after a year of experimental evolution. We addressed the following questions: What genes respond to fluctuating temperature? Does fluctuation promote gene expression evolution? How does the fluctuation regime affect the level of gene expression plasticity?

Using two founder clones we performed a serial transfer experiment for 48 weeks, testing two stable regimes (temperature of 17°C or 23°C), and one fluctuating regime by alternating between these two temperatures every 2.5 days. We identified a pervasive effect of the selection regime on the gene expression level, with a strong difference of evolution between genetic backgrounds. Results show a few hot spots of significantly differentially expressed genes in the genome, in regions enriched with transposable elements (TE), including on dispensable chromosomes. Further, our results evidenced a gain of gene expression robustness associated with the fluctuating regime, suggesting robustness is adaptive in changing environment. Last, an analysis on gene expression correlation revealed a significant set of genes that could act as *trans-*regulators.

## Results

### Mapping quality and normalization

The mapping rate on the genome from the reference strain IPO-323 (Goodwin et al., 2011), was greater than 83.5 % for each sample. Average count of mapped reads was 35 millions after Relative Log Expression (RLE) normalization, and similar normalized read count distributions among samples was obtained (**Additional file 2: Figures S1 and S2**). Correlation coefficient between our sequencing replicates were on average 0.89 (Kendall τ) and agreement between those replicates was fair according to Cohen’s kappa statistic with an average value of 0.2 (McIntyre et al. 2011). Only 10% of the genes changed in expression by more than 2 fold between replicates. We concluded that the repeatability between our experimental replicates was suitable for further analysis (**Additional file 2: Figure S3**). In addition, in comparison to growth at 17°C, growth at 23°C resulted in a greater variation of gene expression (CV = 0.021 *versus* CV = 0.049, at 17°C and 23°C, respectively).

### Genome-wide analysis of the relative abundance of transcripts between acclimated laboratory strains

We checked the quality of our RNA-seq experiment by comparing our data with two previously published studies on the reference strain IPO-323 (Rudd et al., 2015; Kellner et al., 2014). For this purpose we only used the data from the two ancestor clones *MGGP01* and *MGGP44*. Our analysis shows that on average 61% of the genes are transcribed in all samples (with 6663 common gene transcripts on the core chromosomes (CCs) and 39 common gene transcripts on the accessory chromosomes (ACs)) (**Table 1**). FPKM comparisons between pairs of genotypes on CCS show congruent pattern of gene expression (data from 200kb-non overlapping windows, **Additional file 3: Figure S4**). Nevertheless 57% of the common transcripts are differentially expressed between the three genotypes (DESeq2 *Model 1*). Even with a correction for the gene number per chromosome, ACs possess a significantly lower proportion of expressed genes than CCs (*Chi-2* test, *P* < 2.2×10^−16^). Chromosome 7 is singular, presenting significantly less gene transcripts than the other CCs, except for the smallest core chromosome 13 (*Chi-2* tests, *P* < 0.05). Indeed our results confirmed that a large region of 800kb on the chromosome 7 is without any transcriptional activity (**Additional file 3: Figure S5**). This feature was already mentioned for the strain IPO-323 by (Rudd et al. 2015; Schotanus et al. 2015; and Haueisen et al. 2019). This chromosomal region is marked with histone H3K27me3 modifications that mediate transcriptional silencing, and these epigenetic marks are similar to the marks already identified on ACs. This chromosomal fragment is most likely originating from a fusion of a whole accessory chromosome onto the original chromosome 7 (Shotanus et al. 2015). In conclusion, we observed that *in vitro* transcriptome between strains is repeatable for the nature of gene transcripts, but highly variable for the level of expression of gene transcripts.

**Table 1.** Distribution of transcribed genes among chromosomes (Deseq2 model (1)). Number of chromosomes and genes per chromosome for the reference genome IPO-323 (annotation from Grandaubert at al., 2015); total number of genes which are expressed (>4 FPKM) in each of the three genotypes used in this study (*IPO-323*, *MGGP01* and *MGGP44*), and number of common transcripts in all three genotypes; in parentheses: proportion of transcripts among annotated genes; core: chromosomes 1 to 13; accessory: chromosomes 14 to 21.

| IPO-323 annotation |  | Number of expressed genes |  |  |  |
| --- | --- | --- | --- | --- | --- |
| Chromosomes | Gene number | IPO-323 | *MGGP01* | *MGGP44* | Common |
| 1 | 1986 | 1515 (0,76) | 1589 (0,80) | 1604 (0,81) | 1419 (772) |
| 2 | 1146 | 845 (0,74) | 894 (0,78) | 906 (0,79) | 781 (454) |
| 3 | 1073 | 770 (0,72) | 834 (0,78) | 840 (0,78) | 715 (398) |
| 4 | 824 | 605 (0,73) | 652 (0,79) | 667 (0,81) | 554 (324) |
| 5 | 782 | 559 (0,71) | 590 (0,75) | 591 (0,76) | 505 (271) |
| 6 | 692 | 489 (0,71) | 518 (0,75) | 516 (0,75) | 446 (247) |
| 7 | 743 | 385 (0,52) | 402 (0,54) | 412 (0,55) | 347 (190) |
| 8 | 694 | 518 (0,75) | 538 (0,78) | 527 (0,76) | 466 (273) |
| 9 | 604 | 412 (0,68) | 409 (0,68) | 427 (0,71) | 357 (211) |
| 10 | 516 | 359 (0,70) | 375 (0,73) | 376 (0,73) | 325 (189) |
| 11 | 488 | 332 (0,68) | 339 (0,70) | 352 (0,72) | 298 (172) |
| 12 | 414 | 287 (0,69) | 308 (0,74) | 307 (0,74) | 266 (157) |
| 13 | 334 | 210 (0,63) | 230 (0,69) | 224 (0,67) | 184 (110) |
| **Total core** | **10296** | **7286 (0.71)** | **7678 (0.75)** | **7749 (0.75)** | **6663 (3768)** |
| 14 | 114 | 32 (0,28) | 22 (0,19) | 20 (0,18) | 17 (14) |
| 15 | 86 | 14 (0,16) | 25 (0,29) | 8 (0,09) | 6 (4) |
| 16 | 88 | 17 (0,19) | 0 | 4 (0,05) | 0 |
| 17 | 78 | 5 (0,06) | 22 (0,28) | 3 (0,04) | 2 (2) |
| 18 | 64 | 0 * | 18 (0,28) | 0 | 0 |
| 19 | 87 | 18 (0,21) | 14 (0,16) | 11 (0,13) | 7 (6) |
| 20 | 79 | 12 (0,15) | 25 (0,32) | 11 (0,14) | 7 (4) |
| 21 | 58 | 24 (0,41) | 0 | 0 | 0 |
| **Total accessory** | **654** | **122 (0.19)** | **126 (0.19)** | **57 (0.09)** | **39 (30)** |

### Contrasted transcriptomes among evolved lineages

To evaluate whether there was any differences of gene expression between the three selection regimes, we first analyzed the variation of gene expression among all evolved lineages. The ancestral background and the temperature of assay were the two main factors contributing to the total variance (Principal Component Analysis, with 51% and 11.3%, respectively). We also observed a smaller but significant contribution of the selection regimes (6% of total variance) (**Table 2**). We then used *DESeq2* Model *(2)* and found that a large number of genes were significantly differentially expressed (SDE) between the two ancestral backgrounds (7022 genes with an average median fold change of 1.65). We detected 5637 significant genes due to the temperature of assay with an average absolute fold-change of 1.43, and 4146 genes with a significant effect of the selection regime (**Figure 2**). In addition, we detected 1728 SDE genes for the interaction term between ancestral background and selection regime. More specifically, many of those genes showed opposite direction of differential expression between the two ancestral backgrounds (**Additional file 4, Figure S6**). These results indicate that the expression level of a large number of genes evolved differently depending on the genomic context the genes are found. As a consequence, to gain full insight in gene expression changes in response to the selection regime, we considered the two ancestral backgrounds separately for further analysis.

**Figure 1.**
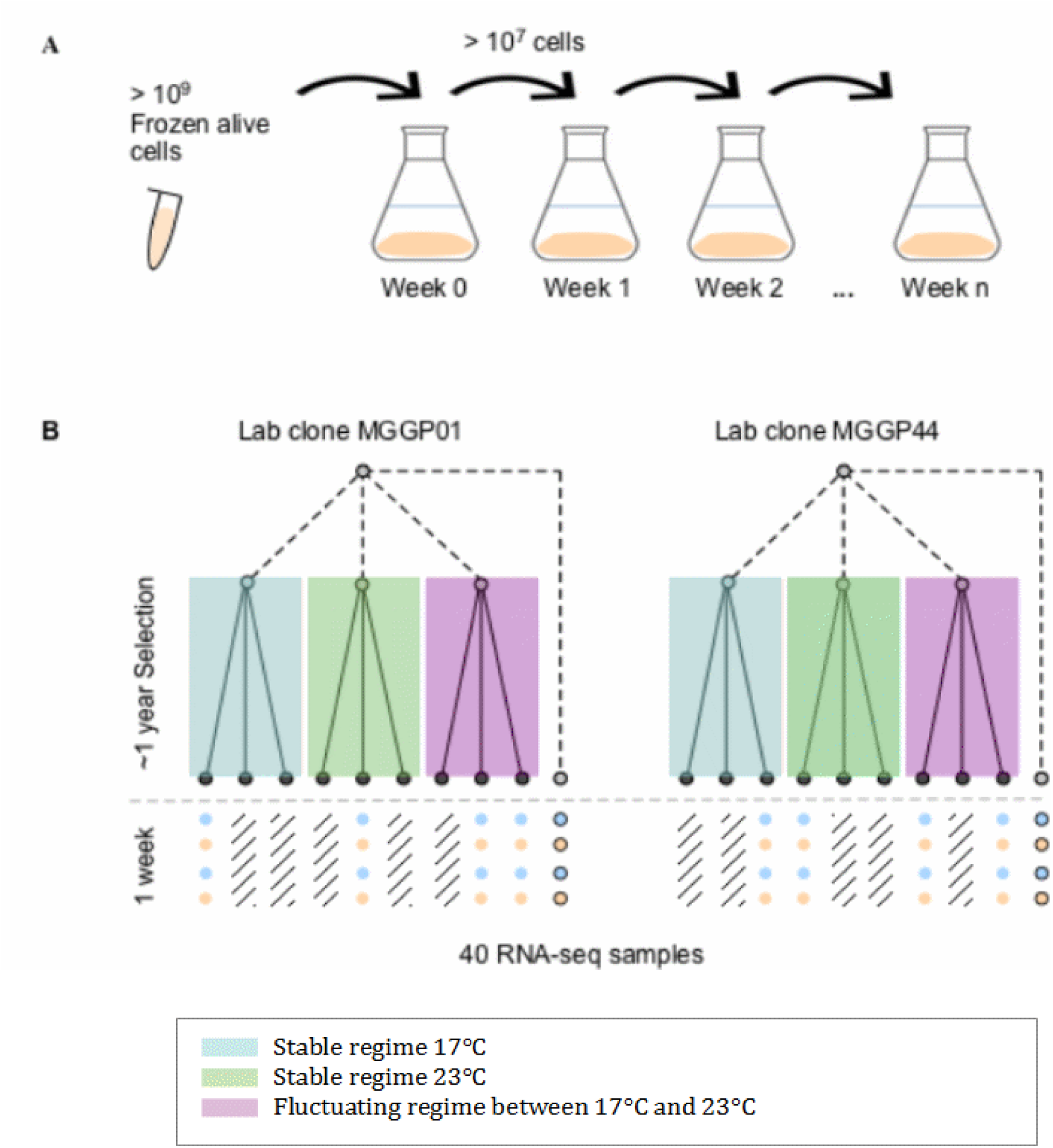
Schema describing the Evolve-and-Resequence approach starting from two lab clones of *Z. tritici*. A: *In vitro* weekly serial transfers occurred over ∼1 year in 500ml liquid medium large erlens; **B:** Description of evolved lineages that were RNA-sequenced for the two ancestral backgrounds, after the experimental evolution under three selection regimes: Stable at 17°C, Stable at 23°C and fluctuating between 17 and 23°C.

**Figure 2.**
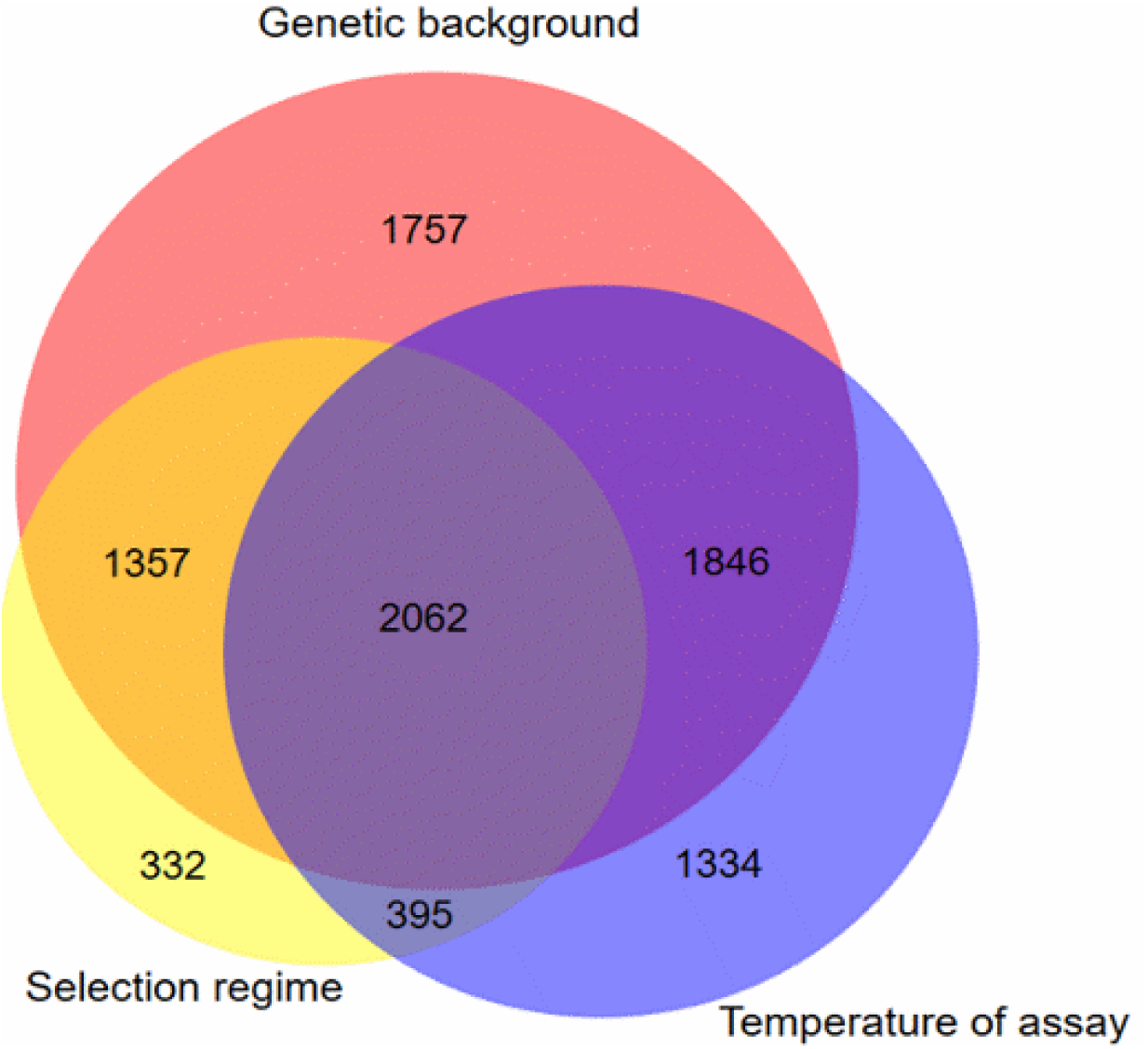
**Venn diagram displaying unique and shared DEGs due to the selection regimes, the temperature of assay and the ancestral genetic background**. Significant genes were obtained from the statistical model DESeq2 model *(2*) with FDR correction for multiple testing correction at 5%.

**Table 2.**
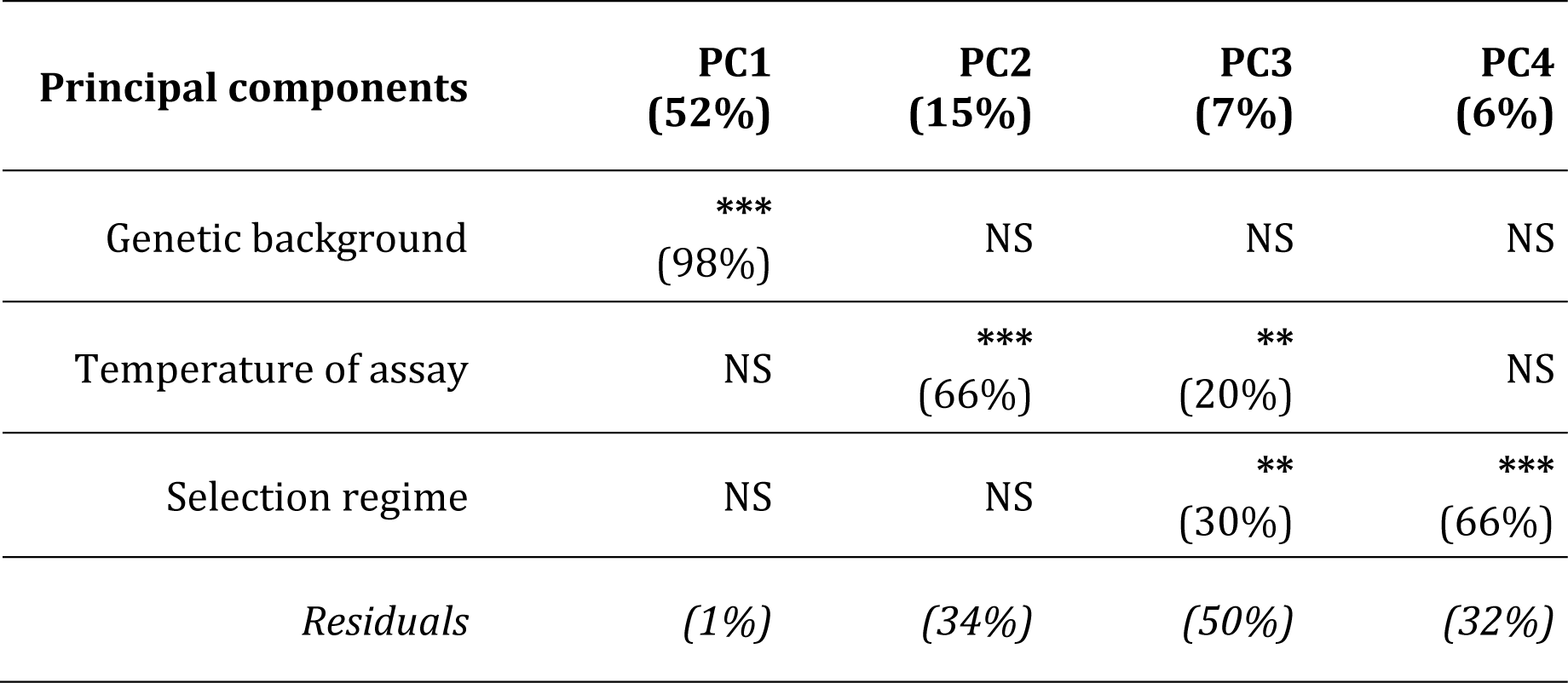
Total percent variance of gene expression explained from DeSeq2 Model (2) (ANOVA). Contribution (percent of variance) of the genetic background; temperature of assay and selection regime to the first four Principal Components (PC1, PC2, PC3, PC4), significance is indicated by NS (non-significant), ** (*P*<0.01) *** (*P*<0.001).

### Evolution of gene expression under the selection regimes

To investigate the effect of the selection regimes on gene expression evolution for each ancestral background, we included the data from the ancestor using our DESeq2 Model *(3)*. To detect evolution of gene expression level in response to fluctuating regime, we seek for genes differentially expressed from the expression level of the ancestor. This analysis is based on two biological independent measurements of gene expression level. On average replicates showed an average Spearman correlation of 0.95, while Cohen’s kappa on FPKM showed a fair agreement of 0.2.

Overall our results indicate that up to 32% of the genes were significantly differentially expressed due to the selection regimes. The temperature of assay also affected the level of expression, for up to 39% of the genes **(Table 3)**. The proportion of genes affected by fluctuation was approximately the same in both ancestral backgrounds (**Table 3, Additional File 5: Figure S7**). The distribution of the fold-change of gene expression for all significant genes is given in the **Additional File 5 (Figure S8)**. Overall, our results seem to suggest that, for each ancestral background, repeated evolution under fluctuation occurred for a significant number of genes (568 and 366 genes, for MGGP01 and MGGP44, respectively) **(Figure 3)**. We focused our attention on this set of genes, hypothesizing those significant changes of gene expression with repeated pattern among experiments could result from adaptive evolution. Unexpectedly, we found an asymmetric response of up- and down-regulated genes between the two ancestral backgrounds, with a majority of down-regulated genes for the background *MGGP01*, while an opposite pattern was found for the background *MGGP44* (**Figure 4**). A large number of those genes were also significant for the stable regimes, to a greater extent for the background MGGP44 (90% and 61% *versus* 21% and 41%, for the regimes Stable 17°C and Stable 23°C, respectively) (**Figure 4, Additional file 5: Figure S9**). While only one single transcript was differentially expressed and specific to the fluctuation regime for the ancestral background MGGP44 (gene ID 58295), 284 genes specifically evolved under fluctuation for the background MGGP01 (**Additional File 6, Table S2**). The contrast between the two genetic backgrounds is strong, illustrating the high genetic diversity level of this fungal species, contributing to the complex genetic basis of gene expression.

**Figure 3.**
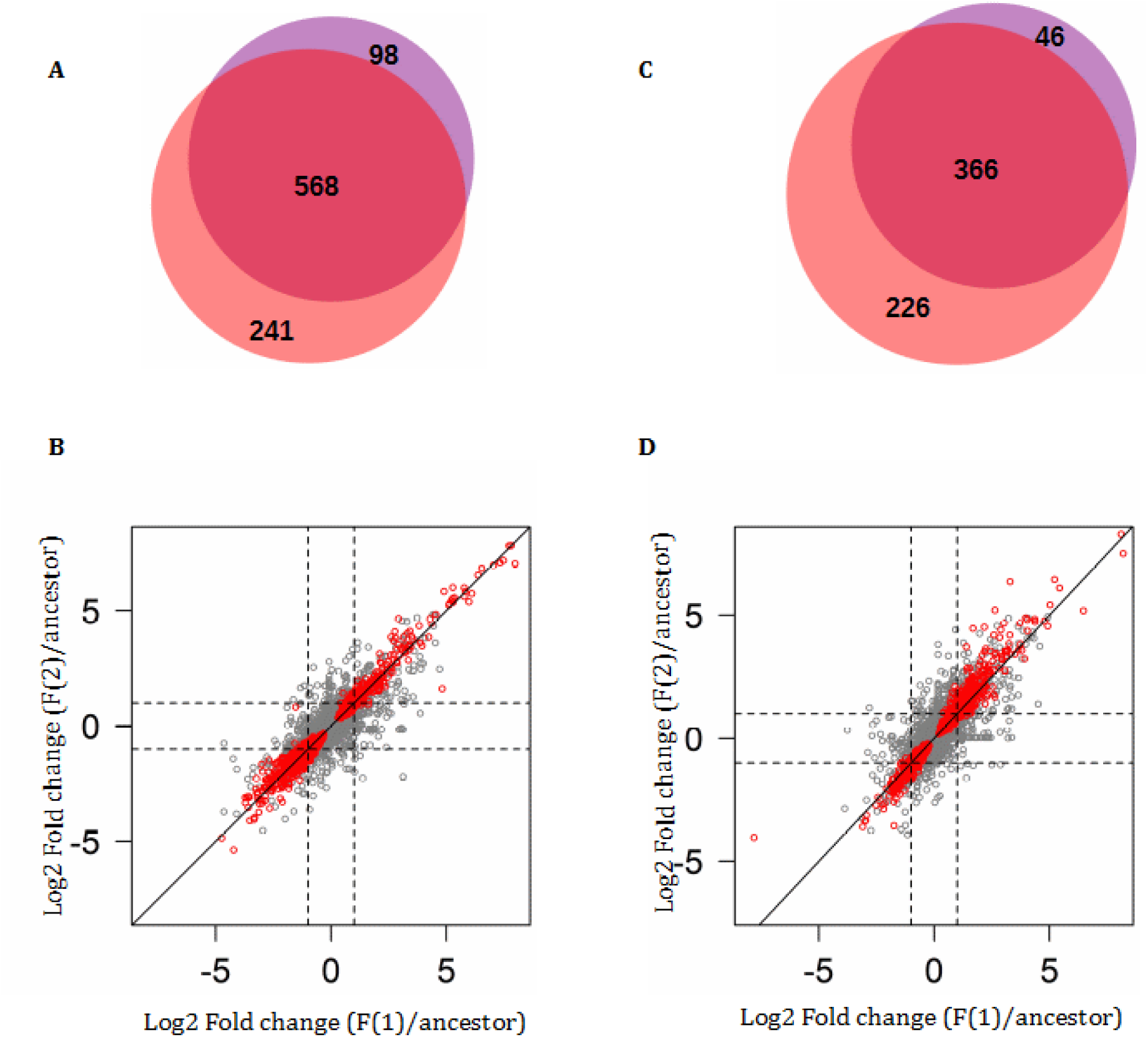
Comparison of significant genes between the two replicates of evolution under fluctuation. **A** and **C**: Venn diagrams showing the number of significant genes which are common between the replicates for the fluctuating regime (**A:** MGGP01, **C:** MGGP44). **B** and **D**: Correlation of Log2 Fold change (evolved lineage against ancestor) between the two replicates under fluctuation (F(1) and F(2)); Pearson’s correlation coefficient on whole data: *ρ =* 0.88 and 0.86, respectively (**B:** MGGP01, **D:** MGGP44). Red points: significant genes after FDR correction at 5%.

**Figure 4.**
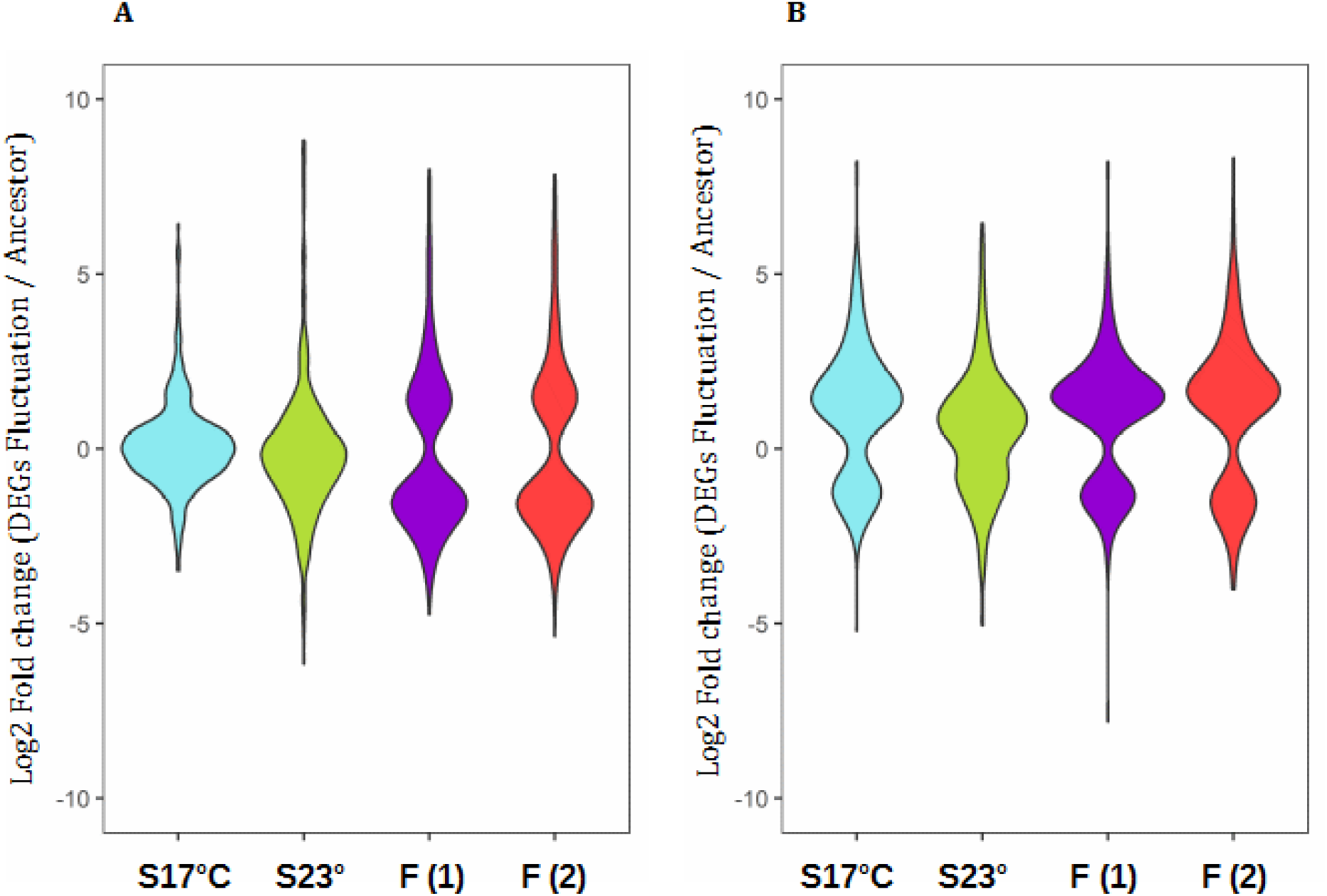
Comparison of gene expression evolution between the stable and fluctuating regimes, for the set of genes with repeated significance in fluctuating regime. Log2 Fold changes are relative to the ancestral gene expression level. **A**: 568 genes for the background MGGP01; **B**: 366 significant genes for the background MGGP44; selection regimes are annotated as follow: Stable regimes (S17°C and S23°C); fluctuating regimes (experimental replicates F(1) and F(2)).

**Table 3.** Number of DEGs from the gene expression level in the ancestor, associated with the selection regime and the temperature of assay. Three selection regimes (stable at 17°C, stable at 23°C and fluctuating between 17°C and 23°C) and two genetic backgrounds (*MGGP01*, *MGGP44*), were tested. Each genetic background was analyzed separately. Significance threshold for genes with a fold-changed >2 was determined after FDR correction for multiple testing at 5% (Benjamini and Hochberg, 2001); median fold change among significant genes when indicated is in parentheses.

|  | Background *MGGP01* | Background *MGGP44* |
| --- | --- | --- |
| Lineage | 3420 | 3659 |
| *Stable 17°C* | 244 (3.1) | 588 (2.9) |
| *Stable 23°C* | 436 (3.1) | 532 (2.8) |
| *Fluctuating* | 666 (2.9) | 412 (2.8) |
| *Fluctuating* | 809 (2.7) | 592 (2.8) |
| Temperature of assay | 3601 | 4333 |
| Lineage-by-temperature | 414 | 851 |

GO term enrichment tests identified different functional terms between the two ancestral backgrounds. First, when considering all genes significant for fluctuation for the background MGGP01 we did not find any evidence for GO term enrichment, while a test including the 284 genes that evolved specifically under thermal fluctuations, detected functional categories related to *Cell cycle control, cell division and chromosome partitioning* (*P* = 0,0019) and *Cytoskeleton* (*P* = 0,027), including proteins known to be essential components of the cytoskeletal mitotic apparatus and essential regulators of cell cycle progression. Notably, the gene length between up- and down-regulated DEGs due to fluctuation was biased toward a smaller size for down-regulated genes (Welch t-tests, *P* = 3.8 × 10-9 and *P* = 0.003, for MGGP01 and MGGP44, respectively). Second, when considering all genes significant for fluctuation for the background MGGP44, significant enrichment for the GO term *defense mechanism*s was found (9 genes, *P* < 0.001).

To fully characterize the transcriptome changes, we examined the distribution of significant genes across the whole genome. First, we reported the number of significant genes among CCs and ACs (**Figure 5A**). For the background MGGP01 only, the proportion of DEGs normalized by the total number of genes per chromosome was higher on ACs than on CCs (**Additional File 7, Table S3**). Second, we surveyed the density of DEGs along each chromosome using a sliding window approach (**Figure 5B and 5C**). The distribution of significant genes under stable and fluctuating regimes was not totally uniform but instead we found 8 regions where the density of significant genes was higher than the nominal threshold of 0.13. Unexpectedly, these hot spots with a higher density –normalized by the number of known annotated genes– of significant genes where often located at the tips of chromosomes: 1, 2 and 6 and 7 for the core chromosomes and 15, 17, 18 and 19 for the accessory chromosomes. To control for a bias due to genomic features, we did the same approach for all significant genes for the temperature effect, and detected no hot spots (data not shown). The presence of significant genes on the accessory chromosomes suggests these chromosomes could be involved in adaptation to abiotic environment. Notice we detected overall 40 genes which expression level was turned off after experimental evolution. None of these genes were located in the hot spot regions, thus excluding the hypothesis that the hot spot regions could simply correspond to deletion events. In addition, 80% of the genes located in the hot spots were up regulated compared to the ancestral gene expression level. We found that the hot spots of DEGs co-locate with regions of high density of TE (TE annotation from Rudd et al. 2015). This might suggest that there is a rapid turn-over of gene regulation in these regions, where TE transposition during mitosis could have an impact on the expression level of the genes.

**Figure 5.**
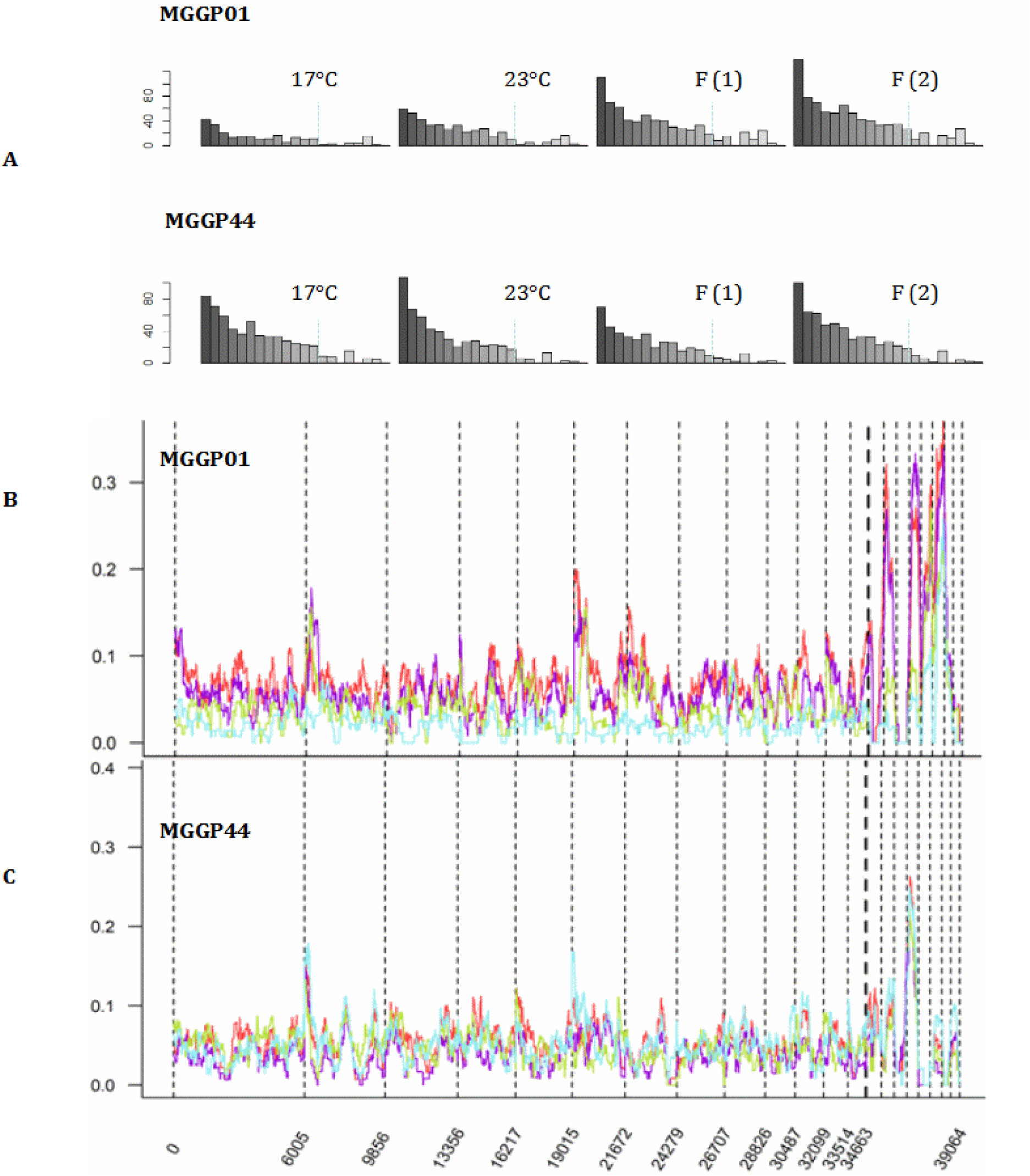
Whole genome distribution of significant genes for a selection regime effect. **A**: Barplots describing the number of significant genes (y-axis) for each of the 21 chromosomes (x-axis), for each condition of experimental evolution tested (Stable: 17°C and 23°C, and replicates under fluctuation: F(1) and F(2), for the two genetic backgrounds (MGGP01 and MGGP44); **B** and **C**: Gene density curves for MGGP01 and MGGP44, respectively. Gene density is given by the number of genes per window of 100kb, using sliding windows with an overlap of 500bp within each chromosome); density curves are presented for each selection regime: Stable 17°C (blue), Stable 23°C (green), fluctuating (1) (purple), and fluctuating (2) (red). Chromosomes are separated by vertical dashed lines.

### Extensive transcriptome rewiring

In the search of sets of genes that would be regulated in concert, we performed a WGCNA analysis. A total of 16 and 15 co-expression modules were detected, respectively for MGGP01 and MGGP44 lineages (**Additional File 8, Tables S4 and S5**). We retained some of the largest co-expression modules that included our most significant genes from our model (*3*) (**Additional File 8, Figure S10**). Consistent with DESeq2 model *(3)* results, this complementary clustering approach largely confirm the most significant enriched categories already identified. We identified the hub gene for each co-expression module. Interestingly, among the 8 selected modules, we also found 24 transcription factors and 39 genes involved in signal transduction with high connectivity **(Additional File 9, Table S6)**. However, based on 10,000 permutation tests to compare the ranking positions of these genes with the ranking positions of 63 randomly chosen genes among the modules, the ranking position of these genes was not significantly biased towards the most connected genes (**Figure 6**). We next investigated the relationship between the physical distance between genes within modules and their level of connectivity. No large *cis*- or *trans-*regulated clusters of genes were detected in our study, but rather scattered co-regulated transcripts all across chromosomes were found on CCs as well as on ACs (**Additional File 10, Figure S11**). Nonetheless our network analysis revealed hub genes and also transcription factors and signal transduction related genes with high connectivity within modules. Whether those genes are genuine *trans*-regulatory factors remains to be demonstrated.

**Figure 6.**
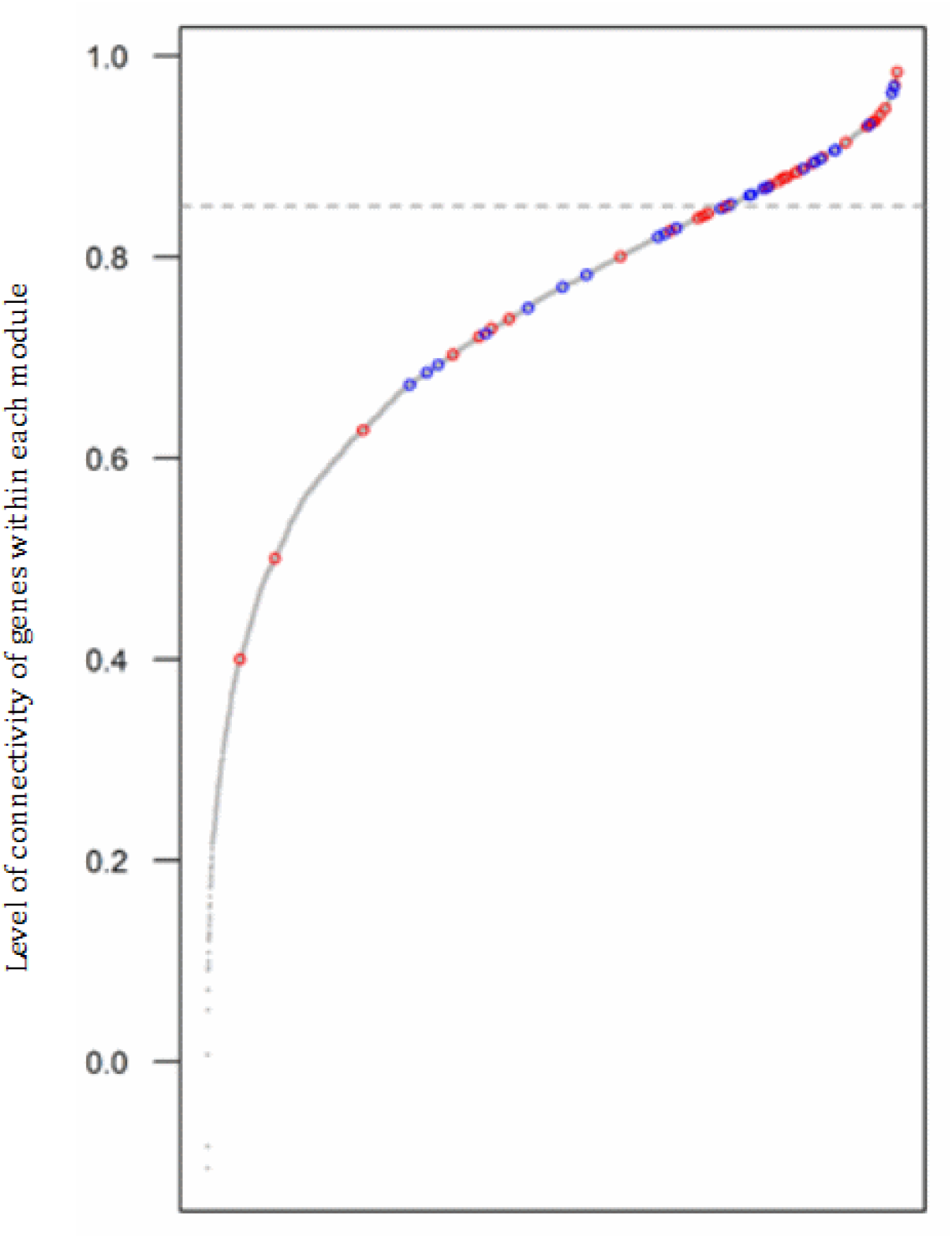
Cumulative distribution of the gene connectivity for all genes included in the. 8 **selected modules revealed by WGCNA.** Gene connectivity is measured by the correlation of each gene to the eigengene within modules. The ranking position of genes with functional annotation related to transcription regulation and signal transduction are indicated in blue and red, respectively.

### Evolution of gene expression plasticity

The effect of the temperature of assay in our laboratory growth conditions —gene expression was assessed at both 17°C and 23°C— was large and pervasive across the genome. We next aimed at determining whether stable or fluctuating temperatures during the EE affected differently the gene expression plasticity. We observed 3.8% and 7.8% of genes significant for Temperature-by-Regime, for MGGP01 and MGGP44, respectively. We also verified that the differential expression between the two temperatures of assay was not due to a change of clone frequency in our evolved lineages. We could directly estimate the presence of new mutations in coding regions and see whether mutations were different between transcripts at the two temperatures of assay. There was no evidence for clonal interference from variants in the coding regions (data not shown). As a consequence a change of transcript level between the two temperatures of assay was assumed to be a plastic change. Using our contrast approach, we could classify genes for gain or loss of plasticity compared to the plasticity of gene expression of the ancestors. We detected a smaller number of genes that became plastic after the experimental evolution for the fluctuating regime (Chi-square tests with Holm’s correction for multiple comparisons, all *P* < 2.10^−7^) (**Figure 7 A and B; Additional File 11, Table S7**). These findings suggest that in our laboratory conditions, evolving under thermal fluctuations promotes robustness. Notably, the relative proportion of plastic genes between the two ancestors is also significantly different (Chi-squared = 33.2, P = 8.10-^9^) (**Figure 7C**). The distribution of the fold-change of gene expression between the two temperatures is biased for both ancestors, but in an opposite manner (with a greater number of up-regulated genes at 17°C as opposed to 23°C, for MGGP01 and MGGP44, respectively).

**Figure 7.**
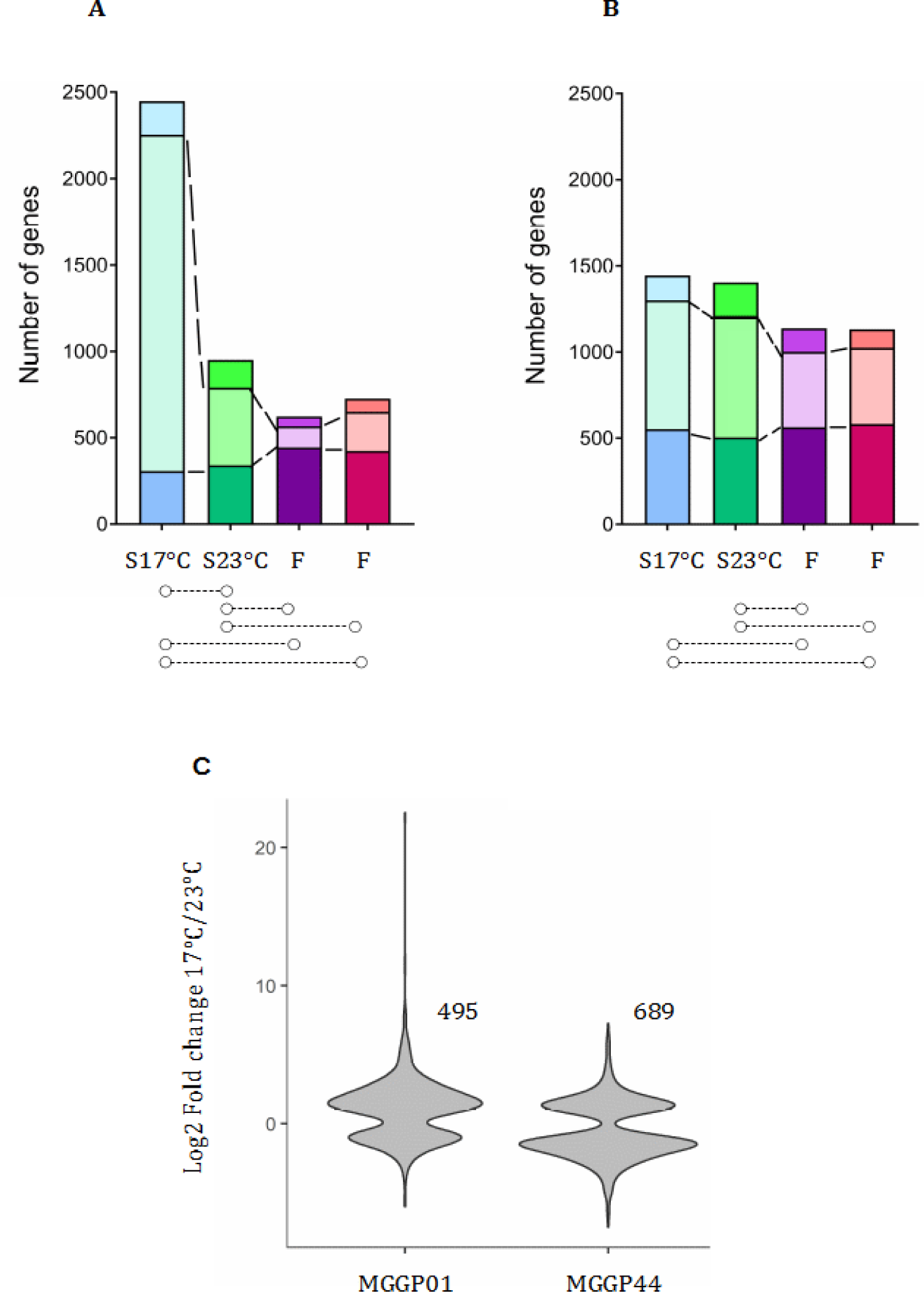
Evolution of plasticity for each selection regime. **A** and **B:** Stacked bar charts describing the number of genes significant for the temperature of assay evolving towards a reduced plasticity (bottom), evolving towards a gain of plasticity (middle bar), and not evolving (top bar). Horizontal dashed lines below the bar charts are indicating the significant pairwise comparisons (Contrast approach using the model DESseq2 *(3)*). **C**: Fold change of gene expression between the two temperatures of assay for the ancestors MGGP01 and MGGP44.

## Discussion

### Intraspecific variation of gene expression

We found pervasive gene expression variation between *Z. tritici* strains, most likely due to the contribution of small-scale mutations (SNPs and small indels). Although comparisons are often made using a limited number of strains, we start to accumulate evidence that there is considerable variation of gene expression within and between populations in *Z. tritici* (Palma-Guerrero et al., 2017; Haueisen et al. 2019). Although high, this amount of variation in *Z. tritici* remains comparable to former results coming from many other studies highlighting the natural variation of transcription within species, mostly using eQTL mapping (*e.g.* Brem, 2002; Schadt et al., 2003; Huang et al., 2015; Zan et al., 2016; Osada et al., 2017).

### Genetic and abiotic factors both contribute to the evolution of gene expression

In this study we quantified the number of genes which expression level is changed due to the EE. Consistent with previous studies, we found a comparable effect of genotype and environment on gene expression level (Hodgins-Davis et al., 2012; Gibson, 2008). Our study based on a ‘moderate’ switch of temperature detects slightly more DEGs due to temperature than due to genetic differences between the founder clones. We also identified a comprehensive set of genes responding to *in vitro* growth and temperature, describing genes and gene functions that can play a primary role in response to abiotic changes in this fungal pathogen. Furthermore, the average cell growth at the two temperatures is not significantly different between ancestors (unpublished data). This observation suggests that generation time, thereby mutation accumulation rate, are likely similar between the two temperatures.

### The effect of fluctuation on transcriptome evolution

Our results suggest that evolution is somewhat repeatable between lineages but show contrasting results between genetic backgrounds. Our results clearly show a pervasive effect of selection regimes on gene expression, with as much as 38% of the *Z. tritici* genome displaying a selection regime effect among evolved lineages. Moreover in our experimental conditions each evolved lineage had on average 530 genes which expression level evolved by at least a factor 2. This result shows a high adaptive potential of the fungal species, which can evolve fast by means of gene expression variation operating on many functional categories. Interestingly, different functional categories were over-represented between the two ancestral backgrounds, suggesting different evolutionary routes can be explored to deal with changes in abiotic environment. The quasi-absence of significant genes in response to fluctuating regime alone for the background MGGP44 is rather surprising. This might suggest that the large number of plastic genes for the ancestor MGGP44 may be advantageous to cope with temperature fluctuations.

### Genome-wide distribution of differentially expressed genes

The genome-wide distribution of all significant DEGs suggests that a few genomic regions are more prone to transcriptional evolution. Compartmentalization of filamentous fungal genomes is a well-known feature, characterized by the presence of regions of distinct evolutionary rate in the genome (Dong et al., 2015, Frantzeskakis et al., 2019). In our experiment, results suggest that regions near subtelomeres on a few core chromosomes and some dispensable chromosomes are more prone to rapid evolution. This result has important implications for the understanding of the capacity of evolution of filamentous fungi. Also this work is the first evidence of rapid adaptation in filamentous fungi, linked to evolution of gene expression on dispensable chromosomes, which are often qualified as a cradle for adaptive evolution, due to their high mutation rates, including structural rearrangements (Croll and McDonald, 2012). The molecular causes of these gene expression changes remained to be understood. As a first attempt towards this goal, we could observe that these regions with higher density of significant genes are also regions enriched in transposable elements. The role of transposable elements in regulating gene expression evolution has been well documented (Sudaran et al., 2014; Rech et al., 2019; Uzunovic et al., 2019; see Schrader and Schmitz, 2019 for recent review). Our results altogether suggest that new mutations –due to TE themselves or SNPS/indels– in the *cis*-regulatory regions of the genes could be responsible for the evolution of gene expression level. In addition, we cannot exclude the hypothesis that some of the cells lost dispensable chromosomes in the evolved lineages, thus leading to a down-regulated effect. This partial loss of accessory chromosomes, if true, would have minor effect since only 20% of the genes under the hot spots are down regulated. Such loss of dispensable chromosome has been observed for 4-week *in vitro* grown cells in yeast-malt-sucrose medium in *Z. tritici* (Moller et al., 2018). The pervasive occurrence of structural rearrangements (indels, translocation, duplications) in filamentous fungi complicates our understanding of the genetic basis of gene expression. To gain full insight into this question, more sequencing of a greater number of evolved lineage and clones within lineages should be conducted.

### Fighting changes by staying stable

We first observed significant differences of plasticity between the ancestors, highlighting genetic variation for transcription plasticity. Genotype-by-environment interaction for gene expression is common in many organisms: *C. elegans* (Li et al., 2006), yeast (Smith and Kruglyak, 2008; Landry et al, 2006; Li and Fay, 2017), *Arabidopsis* genus (He et al., 2016). Here we observed a reduced gain of gene expression plasticity for a significant number of genes for all lineages exposed to fluctuating regime. This result suggests that in our experimental conditions robustness of gene expression is favored under environmental fluctuations. The evolutionary consequences of robustness are important. Robustness can be adaptive and lead to the accumulation of cryptic genetic mutations that can increase the evolvability of the species facing new environments (Le Rouzic and Carlborg, 2008; Cuypers et al., 2017; Payne and Wagner, 2019). Several theoretical approaches have explained a gain of robustness, which is dependent on the rate of fluctuation and the strength of selection (Siegal and Bergman, 2002; LeRouzic et al., 2013). Full understanding of the molecular basis of robustness remains largely unexplored, with a clear lack of empirical support. Main hypothesis proposed that the evolution of robustness under temperature fluctuation is explained by gene networks evolution towards more complexity (Siegal and Bergman, 2002; reviewed in Masel and Siegal, 2009). Robustness could rely on functional redundancy arising from gene duplications, or different network topology. Functional annotation of the genome of *Z. tritici* is not advanced enough yet to gain full insights into these questions. Recently, empirical study in *Arabidopsis thaliana* does not support this hypothesis and demonstrated that a single gene, rather than gene network connectivity or functional redundancy, was leading to trait robustness (Lachowiec et al., 2018). In contrast, another study evidenced robustness by means of functional redundancy in yeast (Li et al., 2010).

### Towards understanding of the genetic basis of gene expression regulation in *Z. tritici*

This work is the first examination the whole transcriptome evolution in response to environmental changes in the filamentous fungus *Z. tritici*. We found widespread variation of up- and down-regulated gene expression in response to temperature, but the genetic basis underlying this variation remains to be explained. Our knowledge on gene expression regulation in this species is little. We cannot exclude in our study that heritable epigenetic variation (histone modification, DNA methylation) are also causing changes in gene regulation. Towards the understanding of the genetic basis of gene regulation in *Z. tritici*, we inferred gene modules based on their co-expression level across lineages. This co-expression network analysis identified 8 hub genes and a significant number of highly connected genes, which are transcription factors, or signal transduction genes. The functional role of those genes remains to be elucidated. Are they genuine capacitors in the gain of robustness?

### Conclusion

Transcriptome evolution analysis at two different temperatures identified a comprehensive set of genes with abiotic variation response and revealed contrasted evolution between the two founder genotypes. These findings illustrate the high potential ability of the fungal species *Z. tritici* to respond to abiotic changes.

## Methods

### Biological material

The two laboratory clones of *Z. tritici MGGP01* and *MGGP44* were collected on wheat in 2010 from the same population sample in the south of France at the location of Auzeville-Tolosane (43° 53’ N – 1° 48’ W). Single colony was isolated and grown in the laboratory using Potato Dextrose Browth (PDB) medium at 17°C, 140 rpm and 70% humidity, for a few weeks prior to the experimental evolution. Strains were then multiplied for storage in PDB medium containing 30% glycerol at −80°C.

### Experimental evolution design

Two clonal strains *MGGP01* and *MGGP44* were used to conduct experimental evolution to test two different genetic backgrounds. Three different selection regimes were tested in triplicates, for a total of 9 experiments starting from each clonal strain. We tested two stable temperatures (17°C and 23°C) and a third condition with fluctuating temperature between 17°C and 23°C every 52-64 hours. Large asexual populations were multiplied in 500 mL of PDB at 140 rpm in a shaking incubator (Infors HT, Inc.). Every week >10^7^–10^9^ cells were transferred into fresh medium by pipetting 20 ml of cell suspension (**Figure 1A**). Large amounts of cell suspension were collected after 48 weeks, to provide liquid stocks at −80°C that were used to produce the RNA samples.

### RNA isolation and sequencing

RNA samples were isolated from the two ancestor clones and 8 evolved lineages (**Figure 1B**). As much samples as possible were sequenced under our budget constraints: 2 fluctuating, 1 stable at 17°C and 1 stable at 23°C for each genetic background corresponding to the lab clones *MGGP01* and *MGGP44.* The 10 samples were grown in the same incubator for one week either at 17°C or at 23°C prior to collect the cell suspensions that were snap-frozen in liquid nitrogen. Each group comprising the 10 samples was grown in duplicate at the two temperatures of assay in separate experiments, in order to create two independent biological replicates Full list of 40 RNA-seq samples is described in **Additional File 1, Table S1**. Prior to RNA extraction of the 40 samples, 50 mL of frozen cell suspension were lyophilized for 3 days (LYOVAC^TM^ GT 2-E freeze dryer, Steris) and ground in liquid nitrogen to a fine powder with a mortar and pestle. Lyophilized tissue was homogenized in 1 mL of Trizol reagent (Invitrogen Inc.), and 200 µL of chloroform, vortexed, incubated at room temperature for 3 min and centrifuged at 12,000 g for 15 min at 4°C. Precipitation of RNA was done by transferring the aqueous phase to a new tube and adding 100% isopropanol. Samples were vigorously shaken and incubated at room temperature for 10 min. RNA pellet was obtained after centrifugation at 12,000 g for 10 min and washed with 1 mL of 70% ethanol. After centrifugation at 7,500 g for 5min, RNA was washed in 70% ethanol, air dried and re-suspended in RNase-free water and stored at −80°C. Sampling quality was checked using agarose gels, Qbit quantification (RNA HS assay kit, Molecular Probes Inc., UK) and Fragment Analyzer (Agilent, CA, USA). Libraries were created using TruSeq Stranded mRNA Sample Prep kit from Illumina by selecting fragments between 250-400bp captured with poly-dT oligonucleotides (Integragen Inc, Evry, France). Paired-end (2 × 75bp) sequencing was done with HiSeq4000 Illumina (Integragen Inc, Evry, France).

### Transcriptome mapping and quantification

We examined raw reads using FastQC (version 0.11.5, (https://www.bioinformatics.babraham.ac.uk/projects/fastqc/), removed adaptors and performed quality-based trimming using Trimmomatic (version 0.32; Bolger et al., 2014) with the following options: PE -phred33 SLIDINGWINDOW:4:20 MINLEN:30 TOPHRED33. For each sample, mate of trimmed reads were mapped to the reference genome IPO-323 (http://genome.jgi.doe.gov/Mycgr3/Mycgr3.home.html) using HISAT2 (version 2.0.4; Pertea et al. 2016). The default parameters were used except for the intron length, the number of multi mapping sites allowed per read and the stringency of mapping score, as follow: -min-intronlen 20 -max-intronlen 15000 -k 3 --score-min L, 0, −0.4. We quantified the number of mapped reads for each predicted gene in the annotation published by (Grandaubert et al., 2015), using Stringtie with the following options: -p 16 -c 7 -m 200 -e –B –G (version 2.3; Pertea et al. 2016). Reads that mapped to the predicted 18S and 28S rRNA genes were all removed from the analysis. The final number of genes utilized for all analyses was 10950.

### Data normalization and differential expression analysis

We performed read count normalization using a size factor calculated for each sample according to the RLE (Relative Log Expression) method implemented in the R package DESeq2 (Love et al., 2014). We used Likelihood Ratio Tests (LRT) followed by Wald tests to compare expression level for each gene. For the comparison of expression levels between genes among genotypes we used FPKM normalization, in order to take in account the gene length.

### Coding variant calling from RNA-sequencing

SNP calling for each RNA-sequencing sample was performed using GATK GenomeAnalysisTK.jar with the following parameters: –T SplitNCigarReads –rf ReassignOneMappingQuality –RMQT 60 –U ALLOW_N_CIGAR_READS and GenomeAnalysisTK.jar –T HaplotypeCaller –dontUseSoftClippedBases –stand_call_conf 20. Custom R scripts were used to identify SNP differences among all samples.

### Natural variation of gene expression level among fungal isolates

Overall gene expression levels were compared between our two ancestor clones and the reference strain IPO-323 (Dutch field strain isolated in 1984), using *in vitro* cultures RNA-seq data from the literature (Kellner et al. 2014; Rudd et al. 2015); (fastq files *SRR1167717*, *SRR1167718* https://www.ncbi.nlm.nih.gov/sra?term=SAMN02640204; and fastq files ERR789217, ERR789218 kindly provided by J. Rudd). Raw fastq files from both studies were analyzed the same way as for our samples.

The following model was fit for each gene to compare the genotypes:

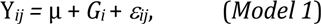

Y*ij* is the expression level of the *i*th genotype (*i* = *IPO-323, MGGP01, MGGP44)* and the *j*th replicate (*j* = 1, 2, 3, 4), µ is the intercept, and ε*ij* is the random error. Significant transcripts were detected by comparing the full model to its null model without the genetic effect, using LRT tests. Correction for multiple testing was performed using a FDR at 5% (Benjamini and Hochberg, 1995).

### Differential gene expression analysis among evolved lineages

To compare the gene expression levels between the evolved lineages the following model was fit for each gene:

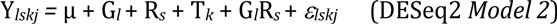

Y*lskj* is the observed expression level of the *l*th genetic background (*l* = MGGP44, MGGP01), the *s*th selection regime (*s* = fluctuating, stable 17, stable 23), the *k*th temperature of assay (k = 17°C, 23°C) and the *jth* replicate (*j* = 1,2), µ is the intercept and ε is the random error. The significance of each term was tested using a LRT by comparing the full model with a reduced model without the considered term. Correction for multiple testing was done using FDR at 5%.

To observe the relative contribution of ancestral background, temperature and selection regimes effects, we also transformed the read counts with the DESeq2 *rlog* function to perform a PCA with the DESeq2 *plotPCA* function. We computed the PC scores of each sample along the first four principal components. Those PC scores were then analyzed with the following ANOVA model:

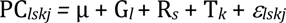

PC*lskj* is the PC score of the *l*th genetic background (*l* = MGGP44, MGGP01), for the *s*th selection regime (*s* = fluctuating, stable 17, stable 23), at the *k*th temperature of assay (k = 17°C, 23°C) and the *j*th replicate (*j* = 1, 2), µ is the intercept and ε is the random error.

### Evolution of gene expression and its relationship with the temperature

For each ancestral background separately we assessed the transcriptome evolution and the level of plasticity of gene expression at the two temperatures of assay. We also tested the interaction between lineages and temperature, which could not be done with the Model (2), due to model over fitting. The following model was fit for each gene and for each ancestral background separately with the R package DESeq2:

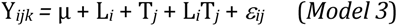

Y*ijk* is the expression level of the *i*th lineage (*i* = *ancestor MGGP01, 12F, 13F, 1217, 1323*, or *ancestor MGGP44, 441F, 443F, 44117, 44323)*, at the *k*th temperature (*k* = 17°C, 23°C), and for the *j*th replicate (*j* = 1, 2), µ is the intercept, and ε*ikj* is the random error. LRT tests were used to compare the full model to reduced models. Wald tests were applied in order to classify significant results using the following rationale:

Among the genes with a significant lineage effect, genes with significant contrast against their ancestor expression level, for both replicated evolution, were considered to have evolved. In addition genes with significant contrast from their ancestral expression level and the expression level for the stable regimes were considered to have specifically evolved under fluctuations.

To identify plastic genes, we considered genes with significant temperature effect, with a fold change greater than 2. To analyze the evolution of plasticity, genes significant for the interaction term temperature-by-lineage were considered using contrasts.

### Gene ontology enrichment tests

Gene ontology enrichment was done using the R package GoSeq (Young et al., 2010). Notice the unknown functional category comprises 5690 genes.

### Gene co-expression network analysis

We search for correlation patterns of expression among genes across our samples. Only genes that displayed at least one significant contrast from our *Model (3)* were included in this analysis (3174 and 3440 genes, for the ancestral background *MGGP01* and *MGGP44*, respectively). Scale-free co-expression networks were built using a Weighted Gene Correlation Network Analysis with the R package WGCNA (weighted correlation network analysis) (Langfelder and Horvath, 2008). Using the log2 transformed read counts, the genes were clustered into modules according to their expression profile in response to the selection regimes and the temperature of assay. Modules were summarized by the first principal component of the gene expression data of the module, and the hub (most highly connected node of the module). Genes within modules were ranked using their correlation to the eigengene of the module.

### Statistical analyses

All statistical tests were performed with R (version 3.3.1, https://www.r-project.org). *Chi*-square tests were used to compare the distribution of the number of significant genes among chromosomes. Welch’s two samples *T*-tests were used to compare the length between up and down-regulated genes which expression level evolved after fluctuation. Chi-square tests with Holm’s or FDR correction (Benjamini and Hochberg, 2001) correction for multiple testing corrections were applied to detect changes of gene expression plasticity. Permutation tests (10,000 bootstraps), were done to test the significance of the ranking position in term of connectivity for the genes with functional annotation associated to transcription regulation or signal transduction.

**List of abbreviations**
AC: Accessory chromosome
CC: Core chromosome
DEG: Differentially expressed gene
FDR: False Discovery Rate
LRT: Likelihood Ratio Test
SNP: Single Nucleotide Polymorphism
TE: Transposable element
WGCNA: Weighted Correlation Network Analysis

## Declarations

### Authors’ contributions

ALR and AG conceived the experiments, AG did the experimental evolution, AJ conducted all the bioinformatics analysis, AJ, ALR, and AG did statistical analyses, interpretation of the data and writing.

## Acknowledgements

We are grateful to Henriette Goyeau for sampling the two fungal isolates and to the genotoul bioinformatics Toulouse Midi-Pyrenees (Bioinfo Genotoul) for providing help and computing resources.

