## Supplemental Table 4, Supplemental Table 5, Supplemental Figure S10 for "Evolution and plasticity of the transcriptome under temperature fluctuations in the fungal plant pathogen *Zymoseptoria tritici*"

### Additional File 8

**Table S4. WGCNA modules (16) identified for the background MGGP01.** Each module contains genes which expression profiles across selection regimes are similar

| WGCNA module | Gene count |
|---|---|
| MGGP01_blue | 760 |
| MGGP01_darkolivegreen | 622 |
| MGGP01_black | 556 |
| MGGP01_magenta | 532 |
| MGGP01_darkgreen | 458 |
| MGGP01_skyblue | 456 |
| MGGP01_red | 356 |
| MGGP01_darkred | 227 |
| MGGP01_purple | 177 |
| MGGP01_tan | 166 |
| MGGP01_orange | 154 |
| MGGP01_darkorange | 151 |
| MGGP01_lightyellow | 125 |
| MGGP01_darkmagenta | 90 |
| MGGP01_white | 77 |
| MGGP01_yellowgreen | 41 |

**Table S5. WGCNA modules (15) identified for the background MGGP44.** Each module contains genes which expression profiles across selection regimes are similar.

| WGCNA module | Gene count |
| --- | --- |
| MGGP44_grey60 | 1509 |
| MGGP44_lightcyan | 1455 |
| MGGP44_black | 409 |
| MGGP44_salmon | 313 |
| MGGP44_green | 304 |
| MGGP44_cyan | 224 |
| MGGP44_darkolivegreen | 185 |
| MGGP44_darkmagenta | 181 |
| MGGP44_tan | 160 |
| MGGP44_darkorange | 144 |
| MGGP44_orange | 101 |
| MGGP44_skyblue | 93 |
| MGGP44_steelblue | 85 |
| MGGP44_violet | 76 |
| MGGP44_sienna3 | 66 |

**Suppl. Figure 10. Characteristics of the top 8 WGCNA modules. A to E:** modules detected for the background MGGP01; **F to H**: modules detected for the background MGGP44. **Top panel** of each figure represents the pattern of the eigengene value among the evolved lineages; **Bottom panel** of each figure represents ranked level of correlation of each gene of the module to the eigengene. Darker points are the ranked positions of genes which are either transcription factors or involved in signal transduction.

**A**

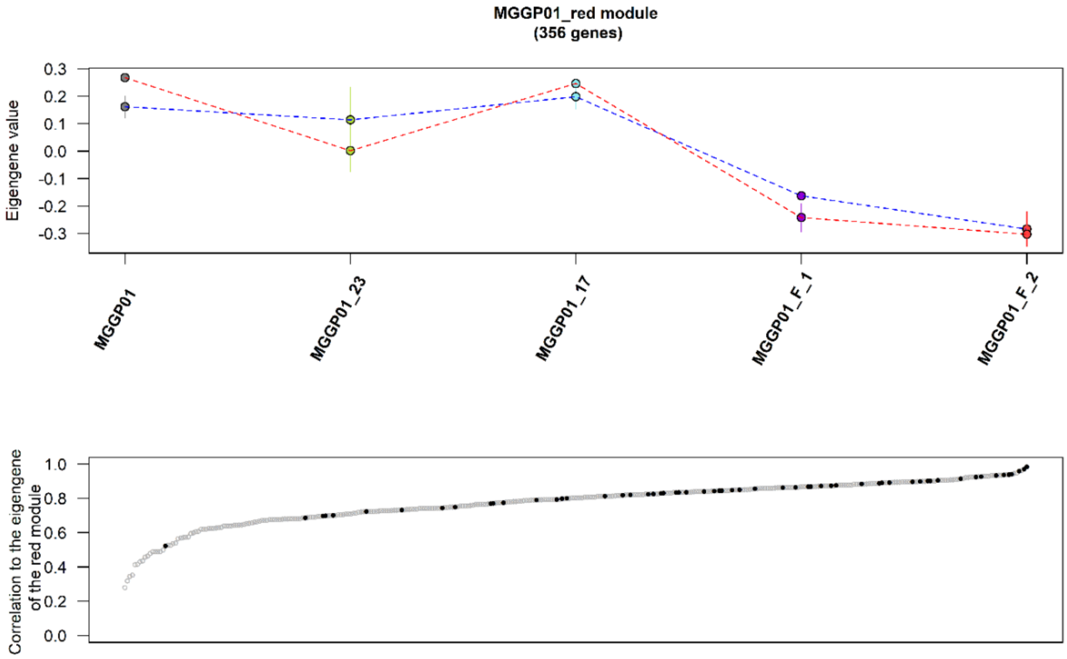

**B**

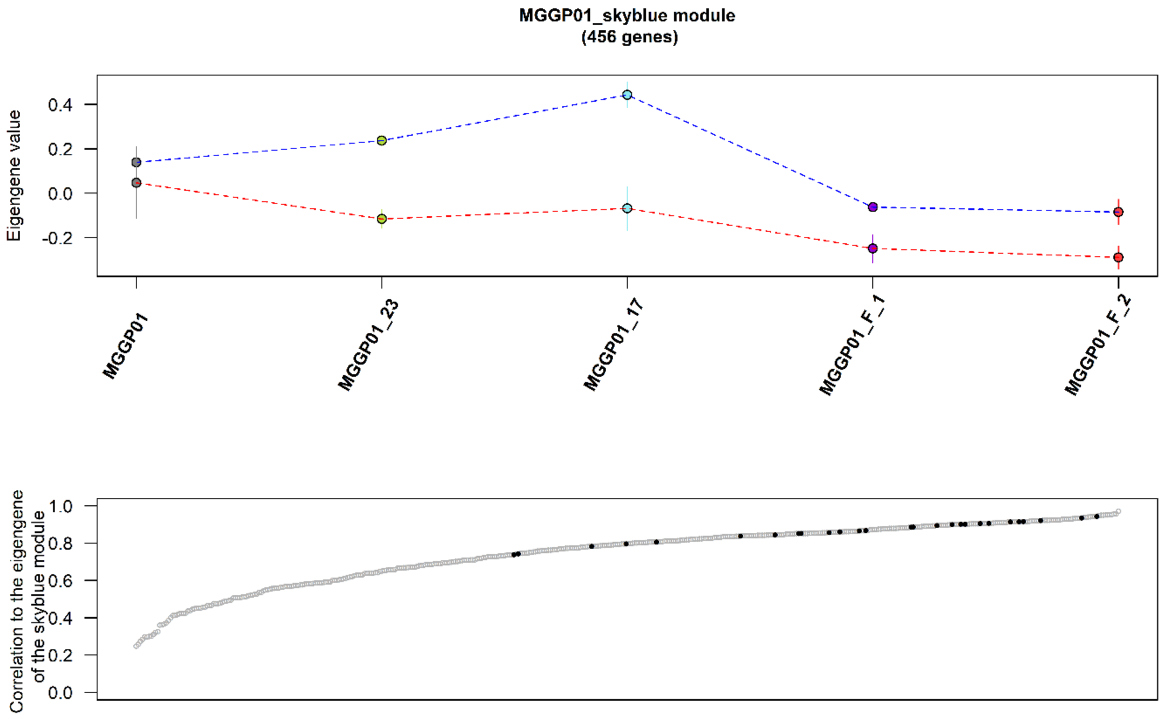
MGGP01_skyblue module
(456 genes)
Eigengene value
0.4
0.2
0.0
-0.2
MGGP01
MGGP01_23
MGGP01_17
MGGP01_F_1
MGGP01_F_2
Correlation to the eigengene of the skyblue module
1.0
0.8
0.6
0.4
0.2
0.0

**C**

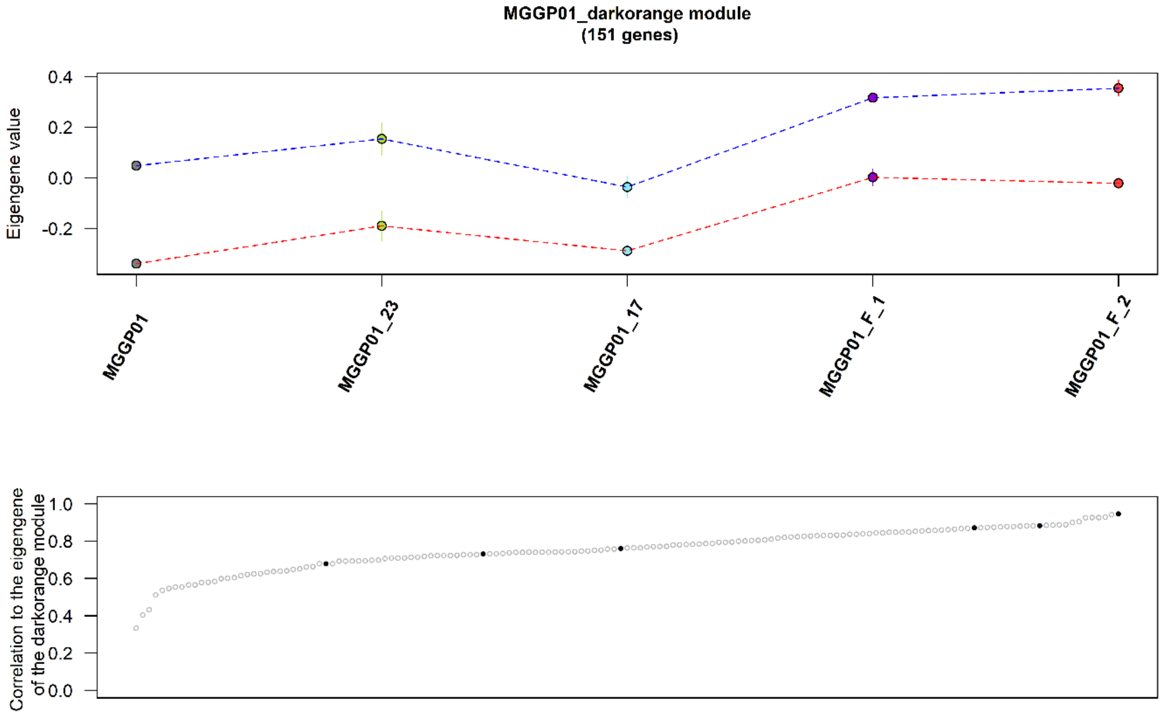
MGGP01_darkorange module
(151 genes)
Eigengene value
0.4
0.2
0.0
-0.2
MGGP01
MGGP01_23
MGGP01_17
MGGP01_F_1
MGGP01_F_2
Correlation to the eigengene of the darkorange module
1.0
0.8
0.6
0.4
0.2
0.0

**D**

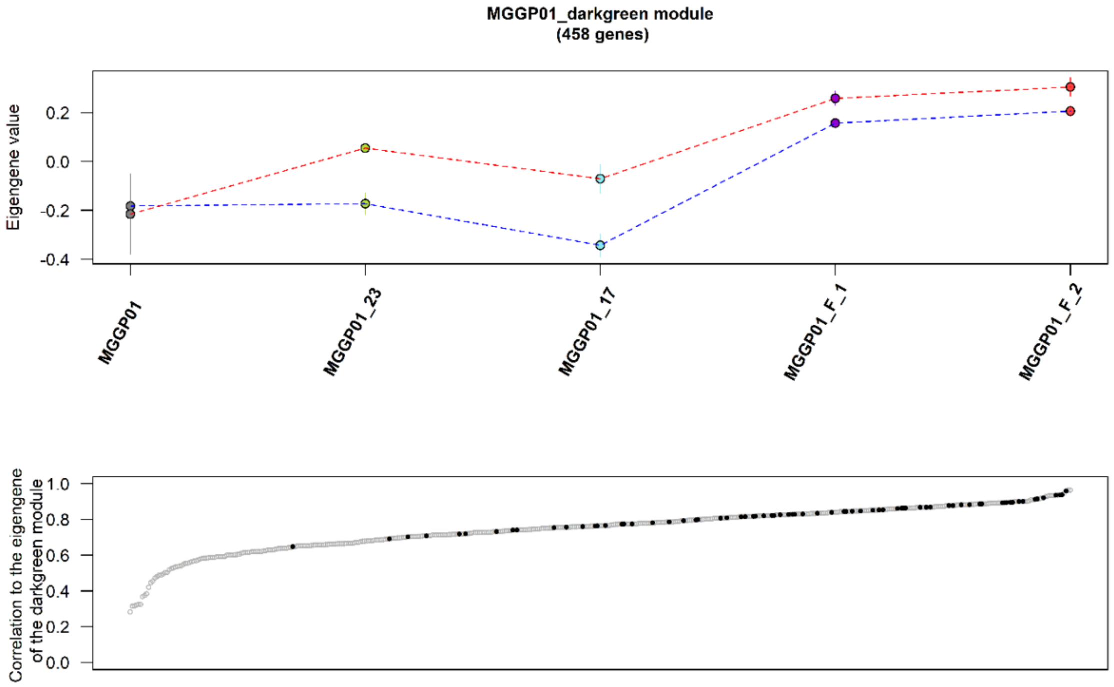
MGGP01_darkgreen module
(458 genes)
Eigengene value
0.2
0.0
-0.2
-0.4
MGGP01
MGGP01_23
MGGP01_17
MGGP01_F_1
MGGP01_F_2
Correlation to the eigengene of the darkgreen module
1.0
0.8
0.6
0.4
0.2
0.0

**E**

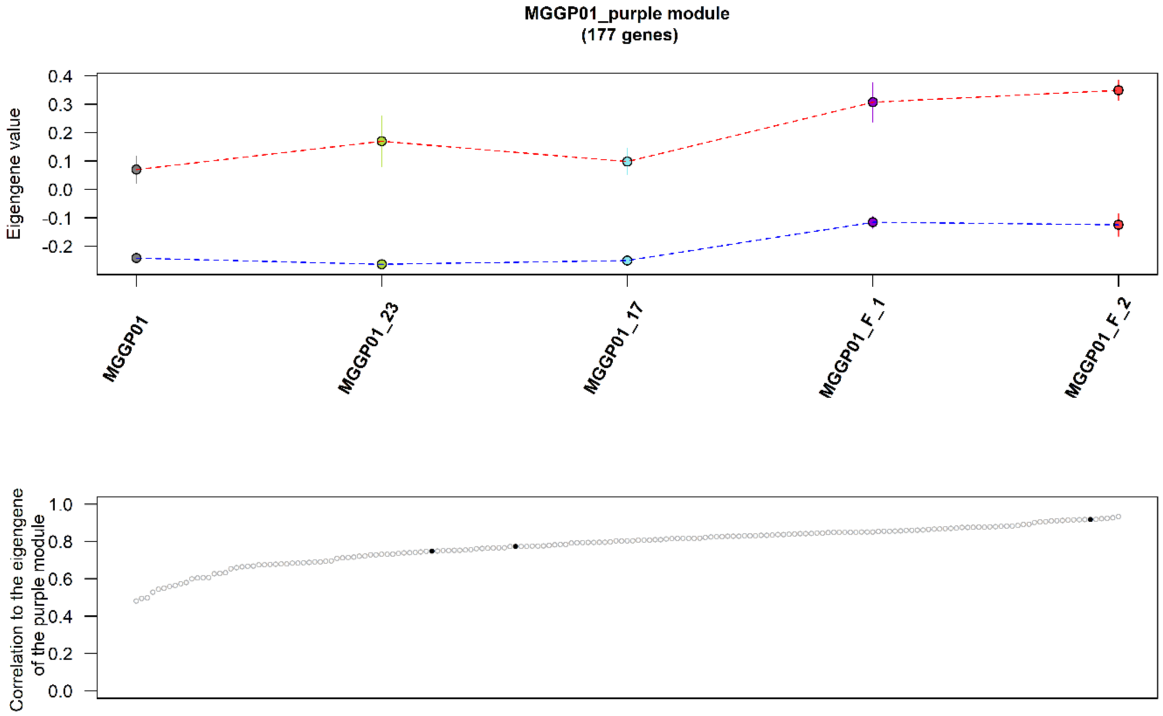
MGGP01_purple module
(177 genes)
Eigengene value
0.4
0.3
0.2
0.1
0.0
-0.1
-0.2
MGGP01
MGGP01_23
MGGP01_17
MGGP01_F_1
MGGP01_F_2
Correlation to the eigengene of the purple module
1.0
0.8
0.6
0.4
0.2
0.0

**F**

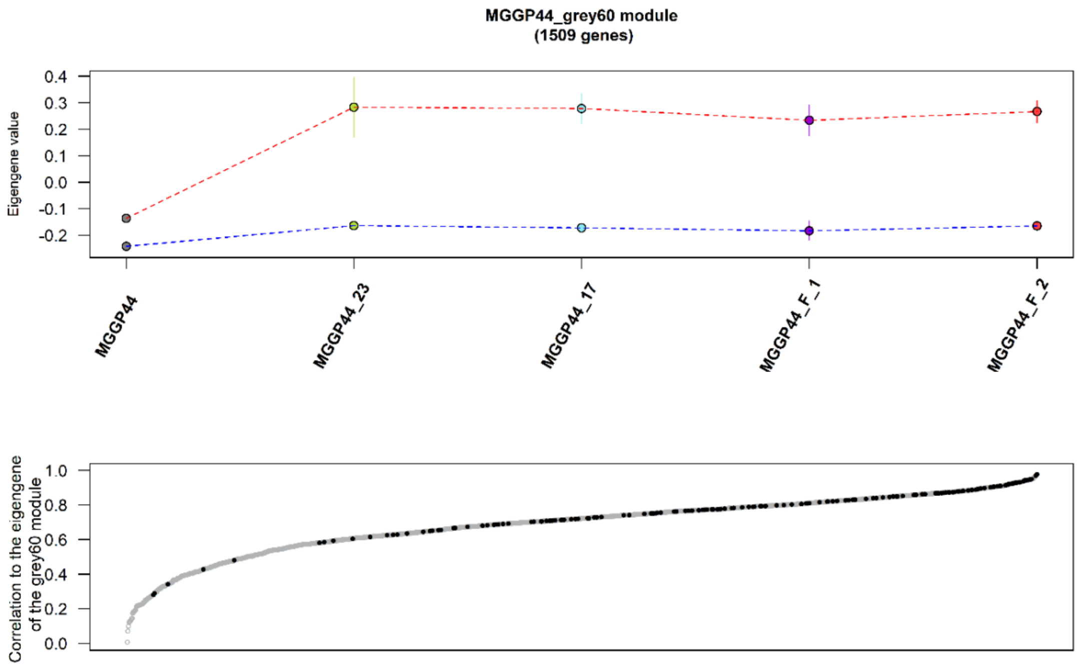
MGGP44_grey60 module
(1509 genes)
Eigengene value
0.4
0.3
0.2
0.1
0.0
-0.1
-0.2
MGGP44
MGGP44_23
MGGP44_17
MGGP44_F_1
MGGP44_F_2
Correlation to the eigengene of the grey60 module
1.0
0.8
0.6
0.4
0.2
0.0

**G**

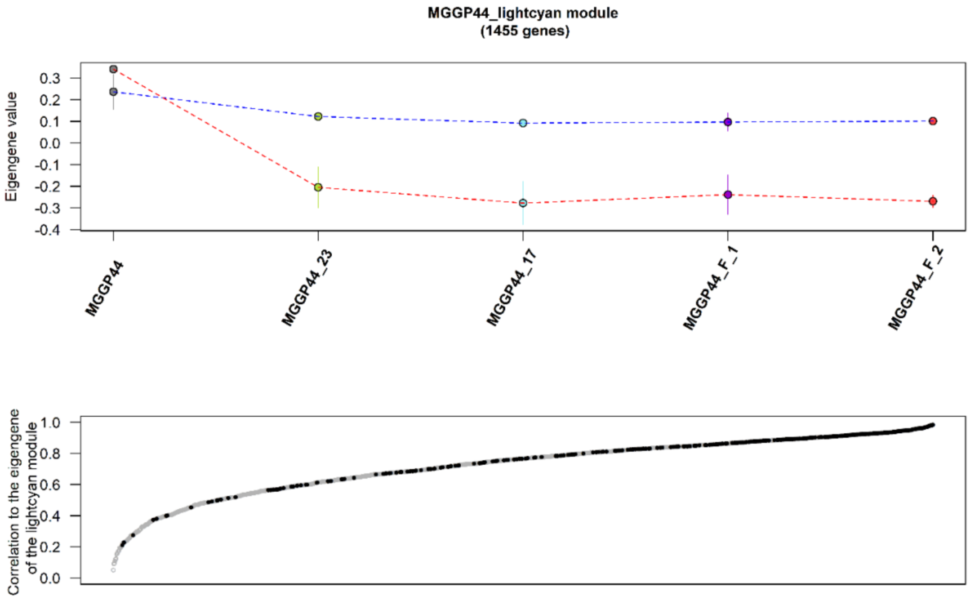
MGGP44_lightcyan module
(1455 genes)
Eigengene value
0.3
0.2
0.1
0.0
-0.1
-0.2
-0.3
-0.4
MGGP44
MGGP44_23
MGGP44_17
MGGP44_F_1
MGGP44_F_2
Correlation to the eigengene of the lightcyan module
1.0
0.8
0.6
0.4
0.2
0.0

**H**

7

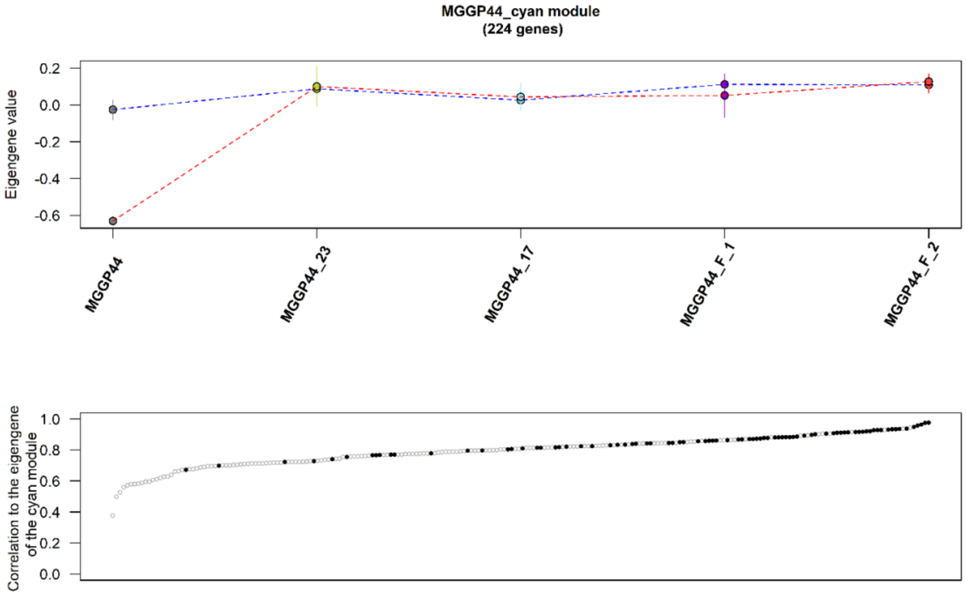
MGGP44_cyan module
(224 genes)
Eigengene value
0.2
0.0
-0.2
-0.4
-0.6
MGGP44
MGGP44_23
MGGP44_17
MGGP44_F_1
MGGP44_F_2
Correlation to the eigengene of the cyan module
1.0
0.8
0.6
0.4
0.2
0.0
