## Supplemental Table 1 for "Evolution and plasticity of the transcriptome under temperature fluctuations in the fungal plant pathogen *Zymoseptoria tritici*"

**Additional File 1: Table S1. List of RNA samples used for the differential expression analysis**

| Nb[1] | Sample IDs | Genetic background | Selection regime | Temperature during selection (°C) | Temperature of assay[2] (°C) | Replicates[3] |
|---|---|---|---|---|---|---|
| 1 | MGGP01_17_R1 | *MGGP01* | - | - | 17 | R1 |
| | MGGP01_17_R2 | *MGGP01* | - | - | 17 | R2 |
| 2 | MGGP01_23_R1 | *MGGP01* | - | - | 23 | R1 |
| | MGGP01_23_R2 | *MGGP01* | - | - | 23 | R2 |
| 3 | MGGP01_23_17_R1 | *MGGP01* | Stable | 23 | 17 | R1 |
| | MGGP01_23_17_R2 | *MGGP01* | Stable | 23 | 17 | R2 |
| 4 | MGGP01_23_23_R1 | *MGGP01* | Stable | 23 | 23 | R1 |
| | MGGP01_23_23_R2 | *MGGP01* | Stable | 23 | 23 | R2 |
| 5 | MGGP01_17_17_R1 | *MGGP01* | Stable | 17 | 17 | R1 |
| | MGGP01_17_17_R2 | *MGGP01* | Stable | 17 | 17 | R2 |
| 6 | MGGP01_17_23_R1 | *MGGP01* | Stable | 17 | 23 | R1 |
| | MGGP01_17_23_R2 | *MGGP01* | Stable | 17 | 23 | R2 |
| 7 | MGGP01_F_1_17_R1 | *MGGP01* | Fluctuating | 17/23 | 17 | R1 |
| | MGGP01_F_1_17_R2 | *MGGP01* | Fluctuating | 17/23 | 17 | R2 |
| 8 | MGGP01_F_1_23_R1 | *MGGP01* | Fluctuating | 17/23 | 23 | R1 |
| | MGGP01_F_1_23_R2 | *MGGP01* | Fluctuating | 17/23 | 23 | R2 |
| 9 | MGGP01_F_2_17_R1 | *MGGP01* | Fluctuating | 17/23 | 17 | R1 |
| | MGGP01_F_2_17_R2 | *MGGP01* | Fluctuating | 17/23 | 17 | R2 |
| 10 | MGGP01_F_2_23_R1 | *MGGP01* | Fluctuating | 17/23 | 23 | R1 |
| | MGGP01_F_2_23_R2 | *MGGP01* | Fluctuating | 17/23 | 23 | R2 |
| 11 | MGGP44_17_R1 | *MGGP44* | - | - | 17 | R1 |
| | MGGP44_17_R2 | *MGGP44* | - | - | 17 | R2 |
| 12 | MGGP44_23_R1 | *MGGP44* | - | - | 23 | R1 |
| | MGGP44_23_R2 | *MGGP44* | - | - | 23 | R2 |
| 13 | MGGP44_23_17_R1 | *MGGP44* | Stable | 23 | 17 | R1 |
| | MGGP44_23_17_R2 | *MGGP44* | Stable | 23 | 17 | R2 |
| 14 | MGGP44_23_23_R1 | *MGGP44* | Stable | 23 | 23 | R1 |
| | MGGP44_23_23_R2 | *MGGP44* | Stable | 23 | 23 | R2 |
| 15 | MGGP44_17_17_R1 | *MGGP44* | Stable | 17 | 17 | R1 |
| | MGGP44_17_17_R2 | *MGGP44* | Stable | 17 | 17 | R2 |
| 16 | MGGP44_17_23_R1 | *MGGP44* | Stable | 17 | 23 | R1 |
| | MGGP44_17_23_R2 | *MGGP44* | Stable | 17 | 23 | R2 |
| 17 | MGGP44_F_1_17_R1 | *MGGP44* | Fluctuating | 17/23 | 17 | R1 |
| | MGGP44_F_1_17_R2 | *MGGP44* | Fluctuating | 17/23 | 17 | R2 |
| 18 | MGGP44_F_1_23_R1 | *MGGP44* | Fluctuating | 17/23 | 23 | R1 |
| | MGGP44_F_1_23_R2 | *MGGP44* | Fluctuating | 17/23 | 23 | R2 |
| 19 | MGGP44_F_2_17_R1 | *MGGP44* | Fluctuating | 17/23 | 17 | R1 |
| | MGGP44_F_2_17_R2 | *MGGP44* | Fluctuating | 17/23 | 17 | R2 |
| 20 | MGGP44_F_2_23_R1 | *MGGP44* | Fluctuating | 17/23 | 23 | R1 |
| | MGGP44_F_2_23_R2 | *MGGP44* | Fluctuating | 17/23 | 23 | R2 |

[1]Nb corresponds to the number of pairs of biological replicates [2]RNA were collected after one week of growth at 17°C or 23°C; [3]two independent biological replicates were produced per lineage
