## Supplemental Figures 1, 2, 3 for "Evolution and plasticity of the transcriptome under temperature fluctuations in the fungal plant pathogen *Zymoseptoria tritici*"

**Additional File 2**

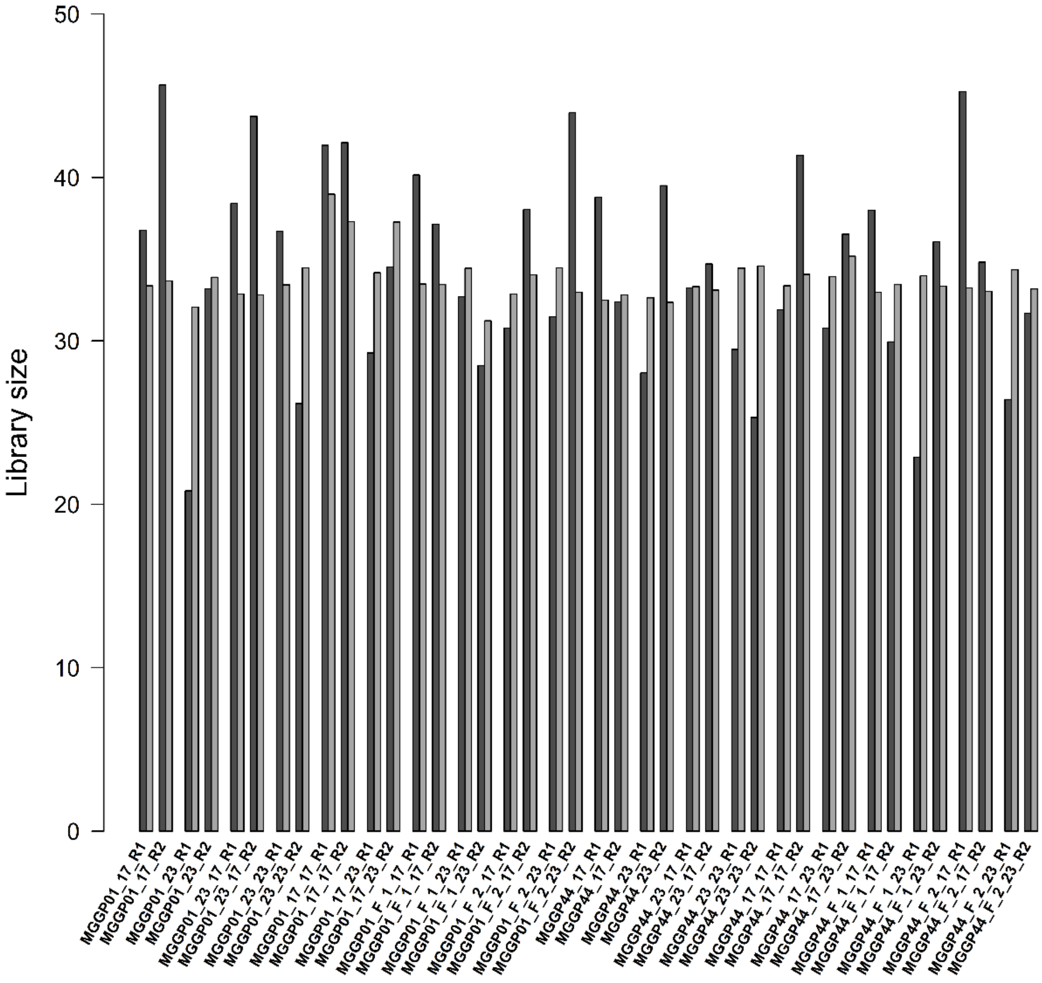


**Figure S1. RNA sample library sizes before and after RLE normalization (y-axis: millions of mapped reads).** Dark grey bars and light grey bars represent library sizes before and after Relative Log Expression normalization, respectively. X-axis labels are described in Table S1. Briefly, two different genetic backgrounds were tested (*MGGP01* and *MGGP44*); 3 different selection regimes were tested (stable 17°C (_17), stable 23°C (_23) and fluctuating temperature between 17°C and 23°C (_F)); 2 experimental replicates for the fluctuating regime were performed (F_1, F_2); 2 temperatures of assay were tested prior to sequencing: 17°C and 23°C (_17, _23); and biological replicates are annotated _R1 and _R2. Reads were mapped using the annotation of the reference genome of *Z. tritici* from (Grandaubert et al., 2015).

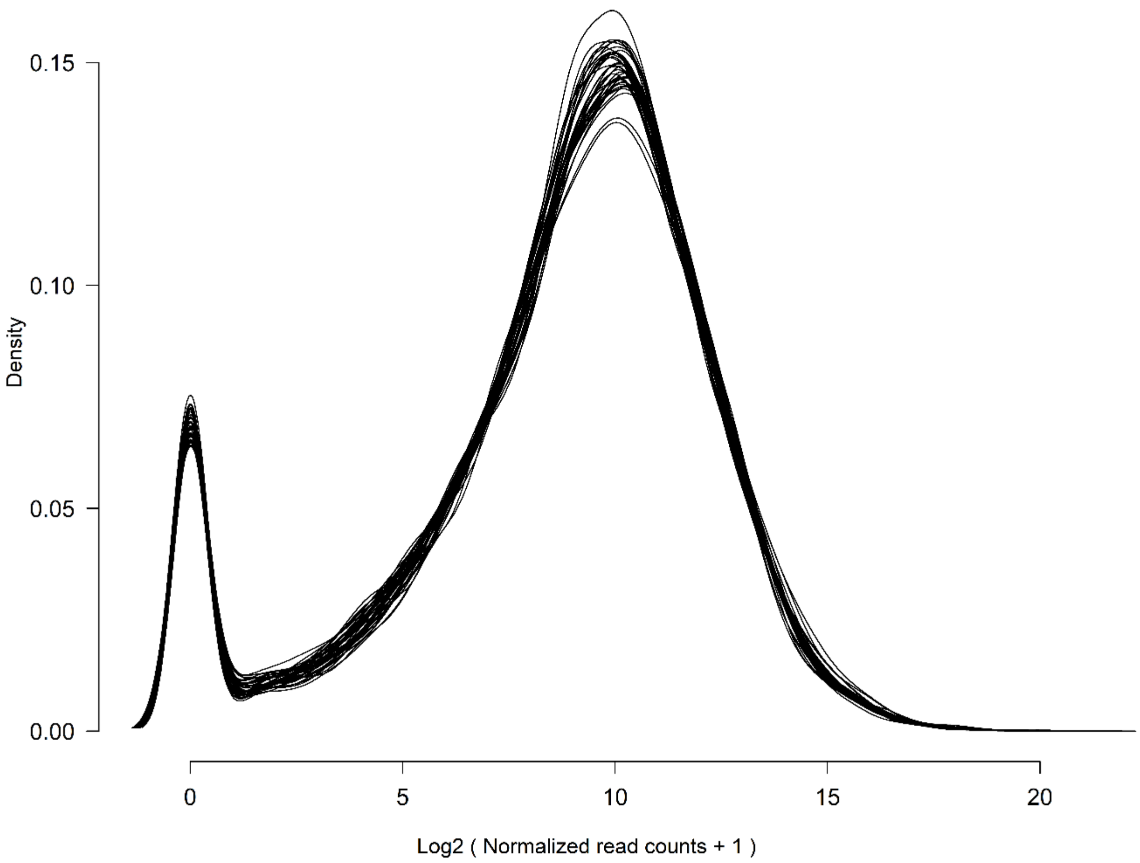


**Figure S2. Distribution of the log2 transformed number of mapped reads per annotated gene for the 40 libraries.**

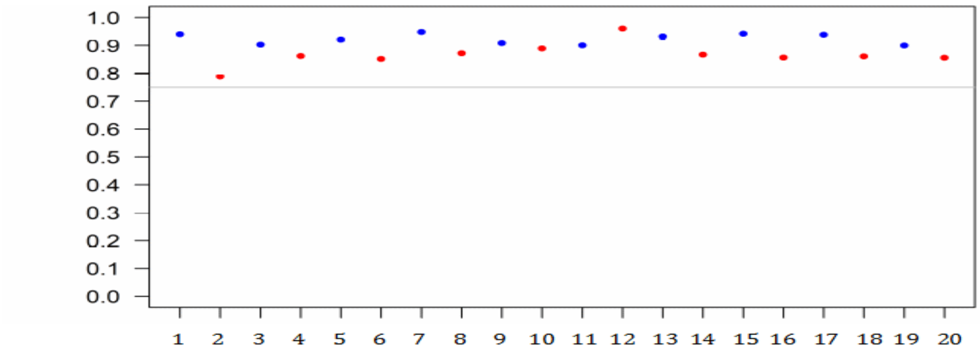


**Figure S3. Kendall's τ correlation coefficient between pairs of biological replicates, estimated using whole transcriptome (10,950 genes annotated in Grandaubert et al., 2015).** Sample numbers and corresponding treatments are listed in the Table S1. Grey line represents the τ value of 0.75.
