## Supplemental Figures 4, 5 for "Evolution and plasticity of the transcriptome under temperature fluctuations in the fungal plant pathogen *Zymoseptoria tritici*"

**Additional File 3**

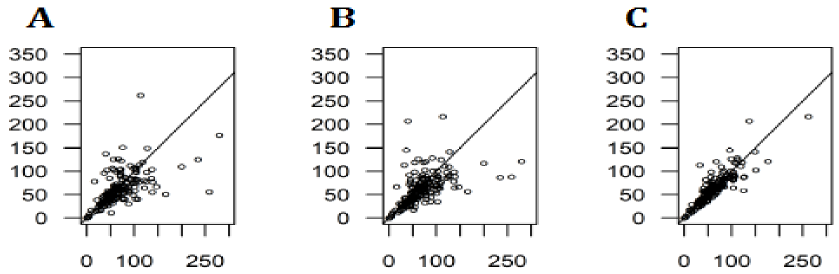


**Figure S4. Scatter plot showing the relationship of averaged FPKM between pairs of genotypes. A**. Strains *IPO-323* and *MGGP01***; B.** *Strains IPO-323* and *MGGP01*; **C.** *MGGP01* and *MGGP44*. 181 non-overlapping windows of 200 kb were used. Pearson's correlation coefficients are 0.55, 0.52 and 0.76, for A, B and C, respectively; and the black line corresponds to the first bisector (y = x).

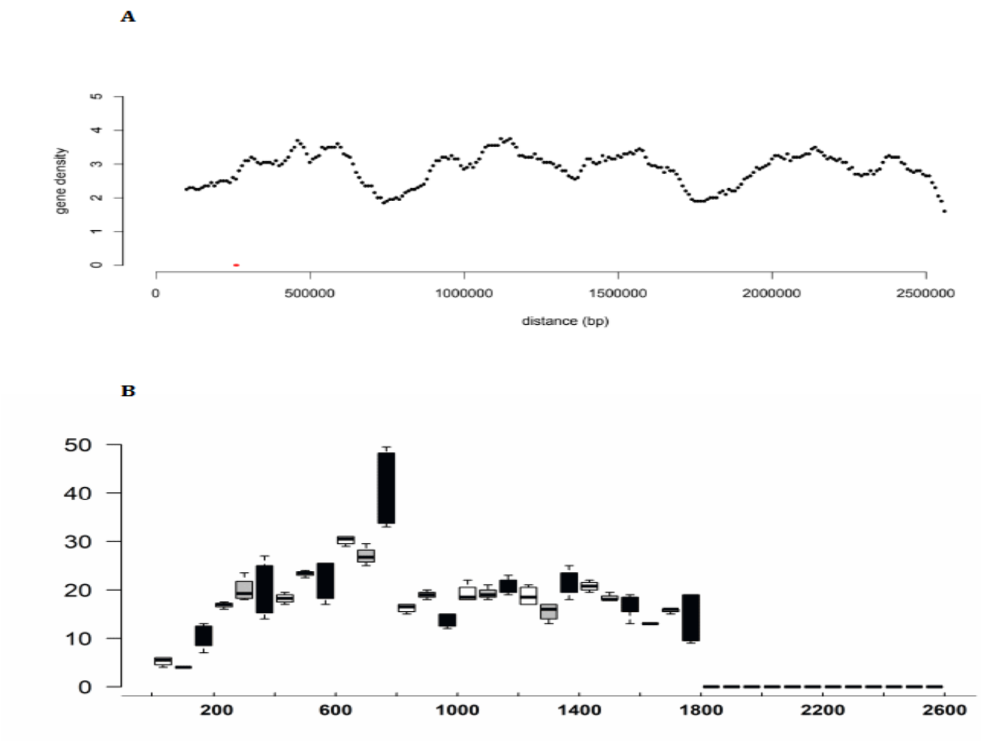


**Figure S5. Transcriptional activity on chromosome 7**

**A.** Gene density along the chromosome 7 (30Kb overlapping windows by 10Kb); **B.** Gene expression profile for three isolates *MGGP01* (white), *MGGP44* (grey), and *IPO-323* (black), using median FPKM within 200kb non-overlapping windows for all annotated genes along the chromosome 7 (Grandaubert et al., 2015).
