## Supplemental Figure 6 for "Evolution and plasticity of the transcriptome under temperature fluctuations in the fungal plant pathogen *Zymoseptoria tritici*"

**Additional File 4**

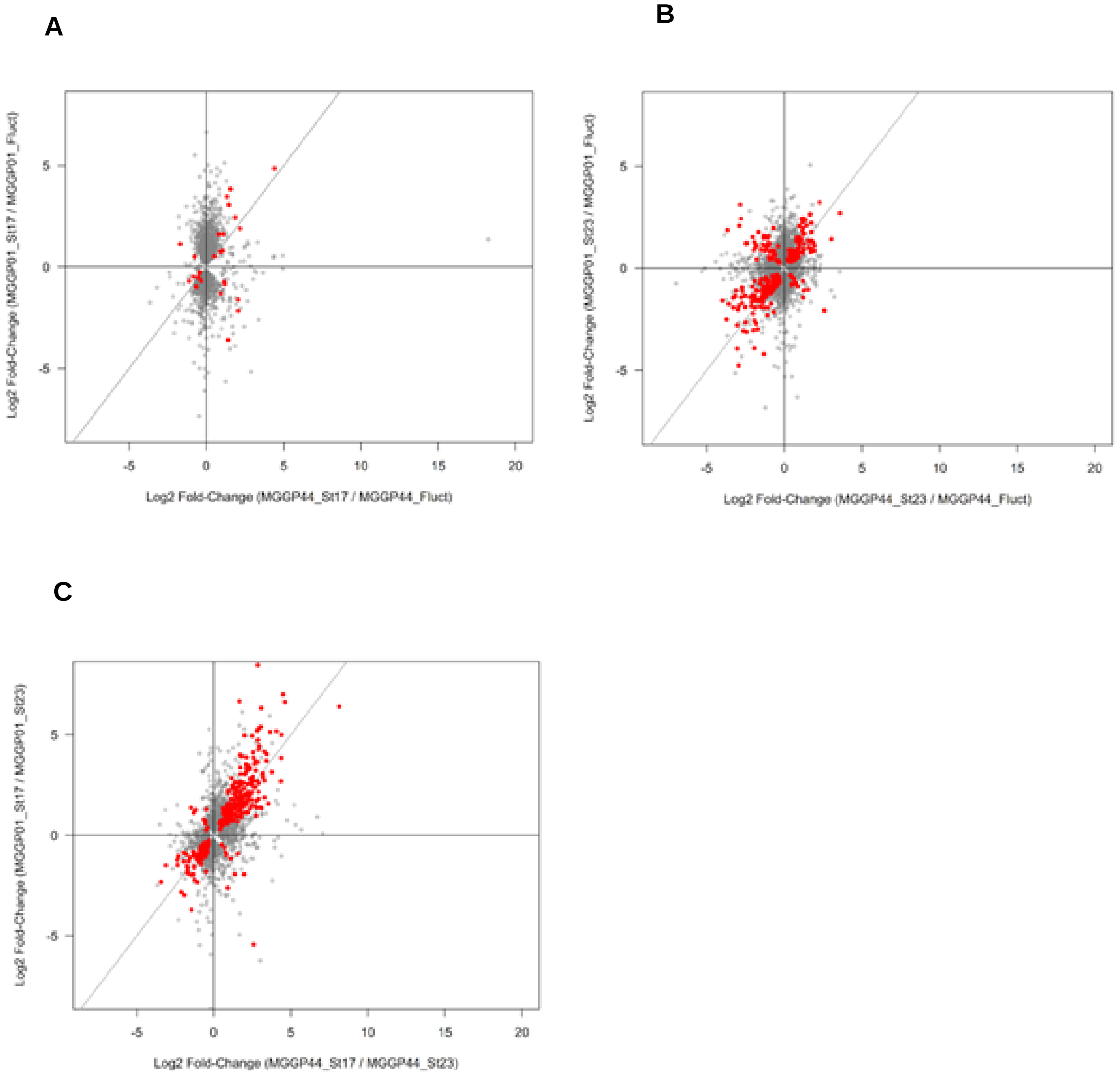


**Figure S6. Correlation between fold changes of gene expression among selective regimes measured for the two genetic backgrounds MGGP44 and MGGP01.** All significant genes (1728) for the interaction term Regime-by-Genetic background in model DESeq2 (2) are included; Red points: significant genes between the selection regimes using a contrast approach; **A:** Fold change of gene expression between Stable at 17°C and Fluctuating lineages, Spearman correlation coefficient $\rho = 0.23$; **B:** Fold change of gene expression between Stable at 23°C and Fluctuating lineages, Spearman correlation coefficient $\rho = 0.31$; **C:** Fold change of gene expression between Stable at 17°C and Stable 23°C lineages, Spearman correlation coefficient $\rho = 0.51$.
