## Supplemental Figures 7,8,9 for "Evolution and plasticity of the transcriptome under temperature fluctuations in the fungal plant pathogen *Zymoseptoria tritici*"

**Additional File 5**

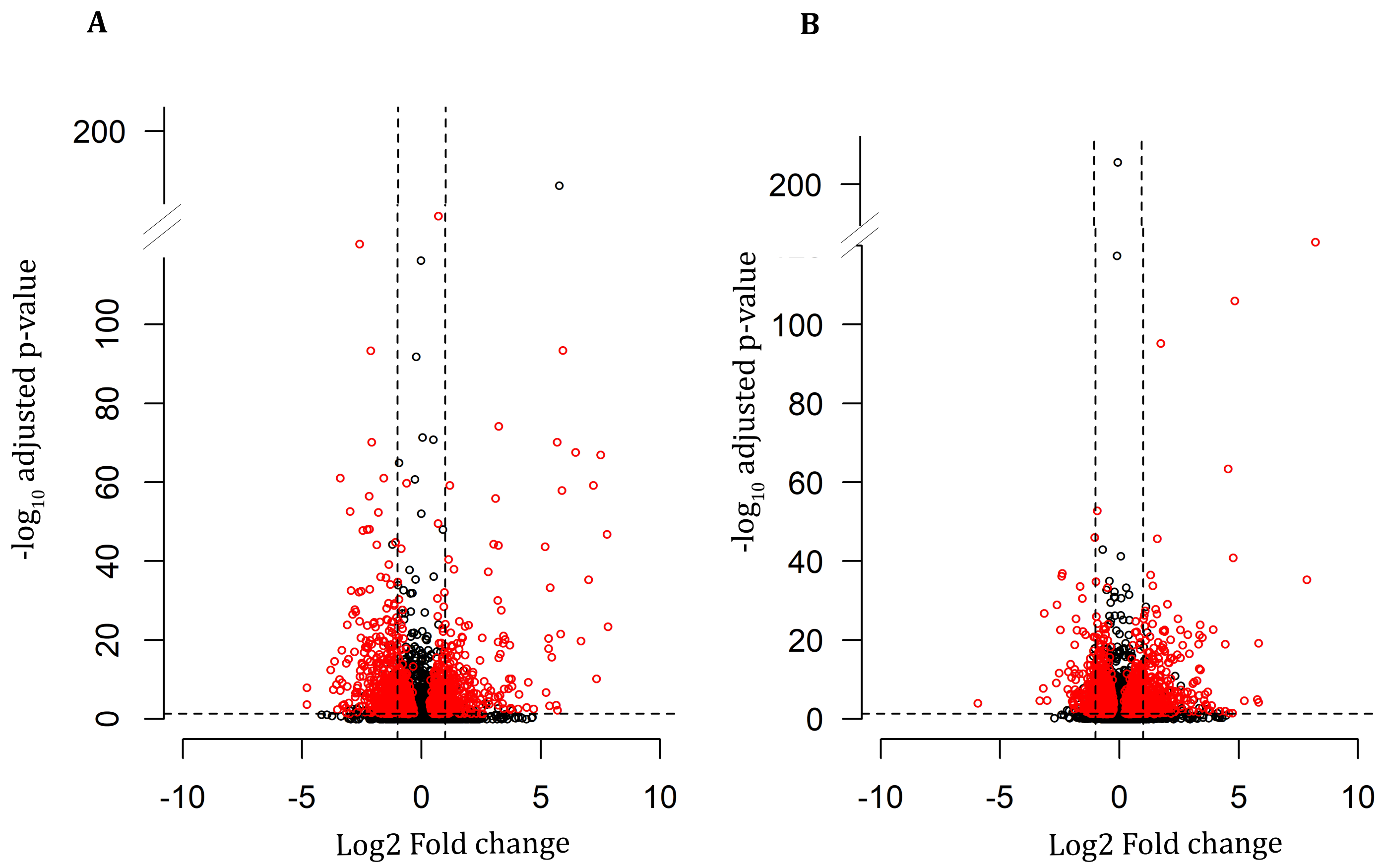


**Figure S7. Volcano plots of DESeq2 analysis model *(3)* for DEGs compared to the ancestor expression level. A:** results for the genetic background MGGP01 ; **B:** results for the genetic background MGGP44 ; vertical dashed lines: Fold change of 2 ; horizontal dashed line: FDR threshold at 5 %.

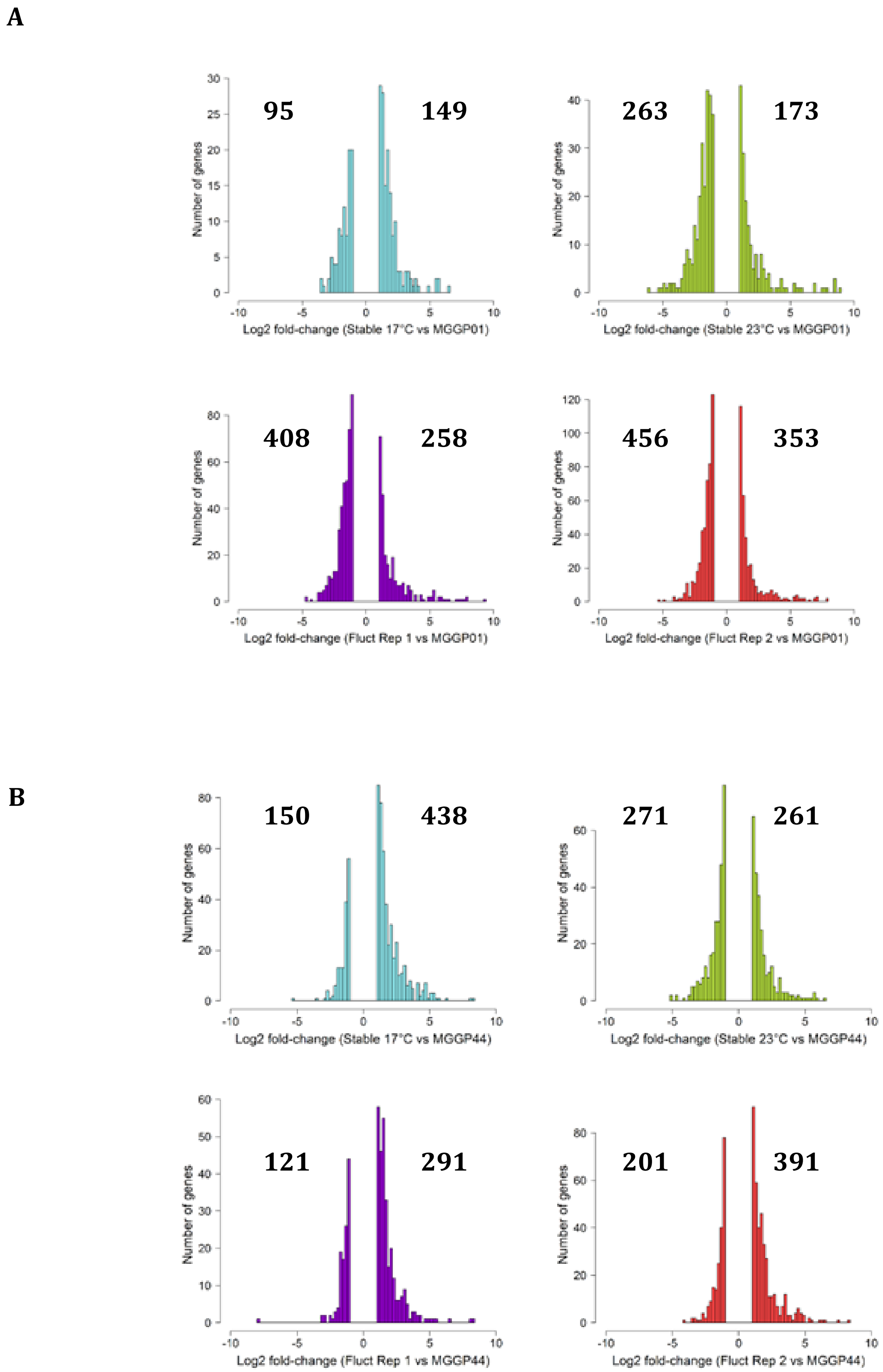


**Figure S8. Distribution of Log2 fold change of all significant genes differentially expressed due to the selection regimes in the model DESeq2 *(3)*. A:** results using the genetic background MGGP01 ; **B:** results using the genetic background MGGP44; number of up- and down-regulated significant genes are indicated on top of each graph.

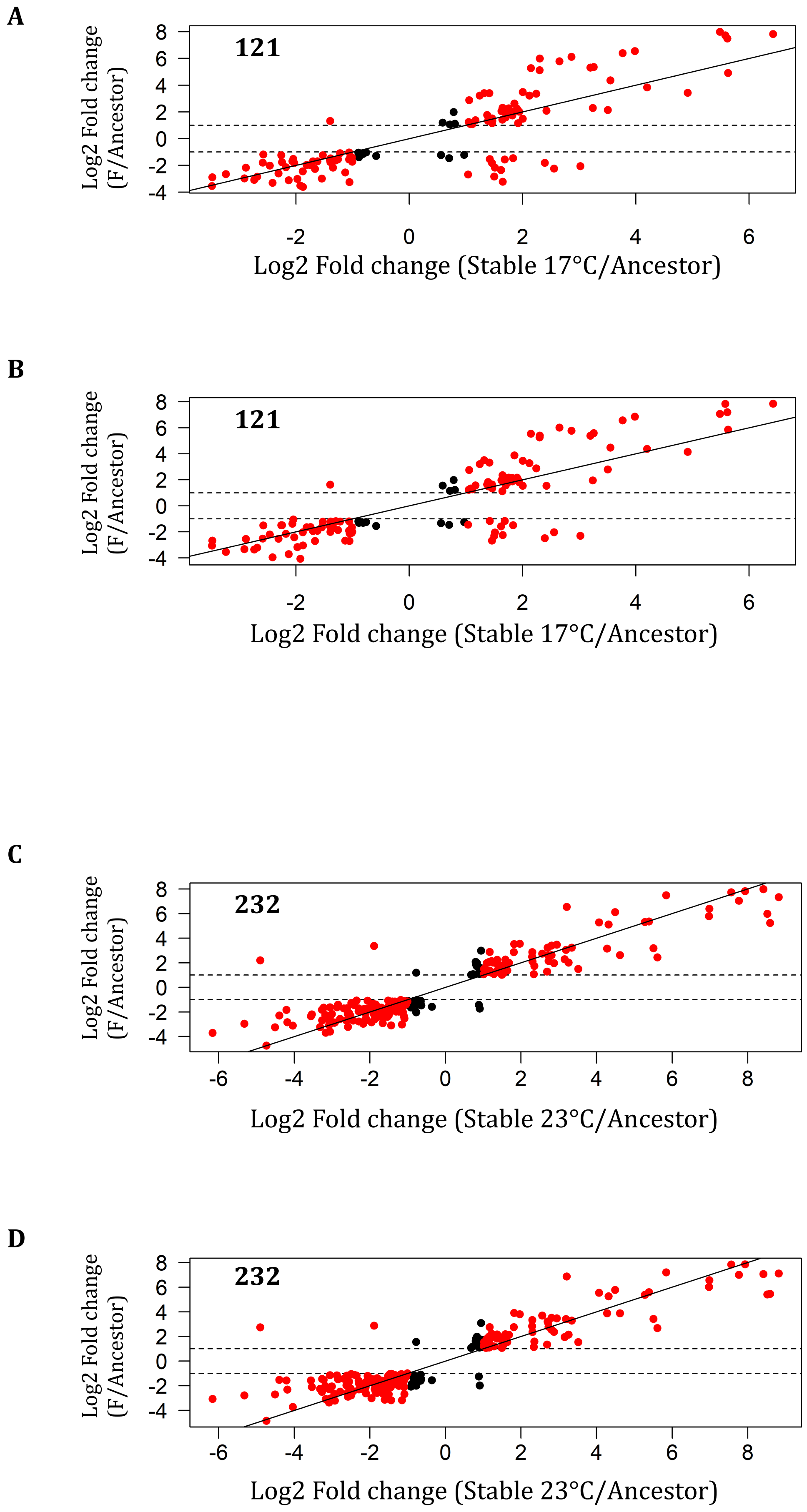
A
121
Log2 Fold change (F/Ancestor)
Log2 Fold change (Stable 17°C/Ancestor)
B
121
Log2 Fold change (F/Ancestor)
Log2 Fold change (Stable 17°C/Ancestor)
C
232
Log2 Fold change (F/Ancestor)
Log2 Fold change (Stable 23°C/Ancestor)
D
232
Log2 Fold change (F/Ancestor)
Log2 Fold change (Stable 23°C/Ancestor)

**E**

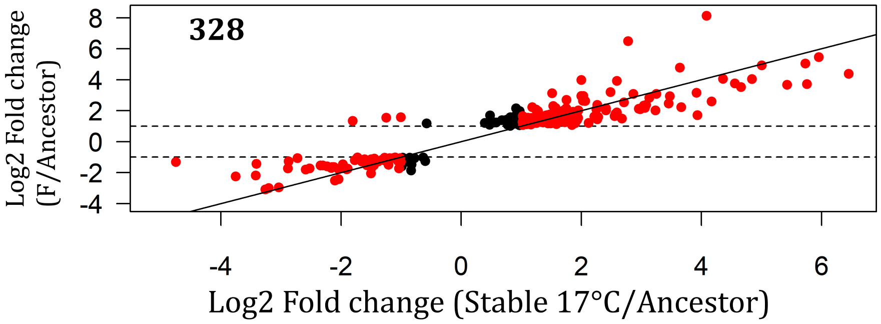
328
Log2 Fold change (F/Ancestor)
Log2 Fold change (Stable 17°C/Ancestor)
8
6
4
2
0
-2
-4
-4
-2
0
2
4
6

**F**

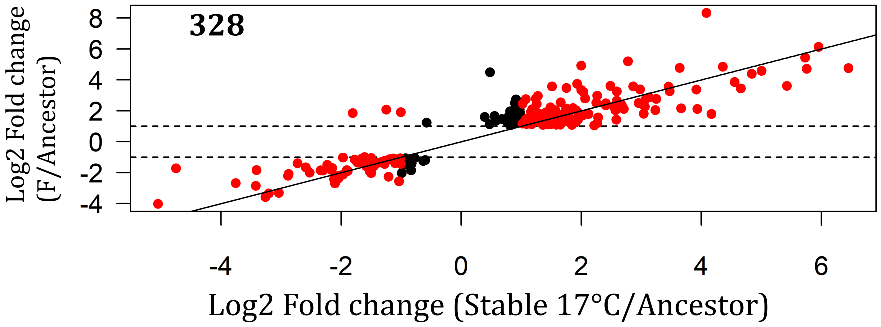
328
Log2 Fold change (F/Ancestor)
Log2 Fold change (Stable 17°C/Ancestor)
8
6
4
2
0
-2
-4
-4
-2
0
2
4
6

**G**

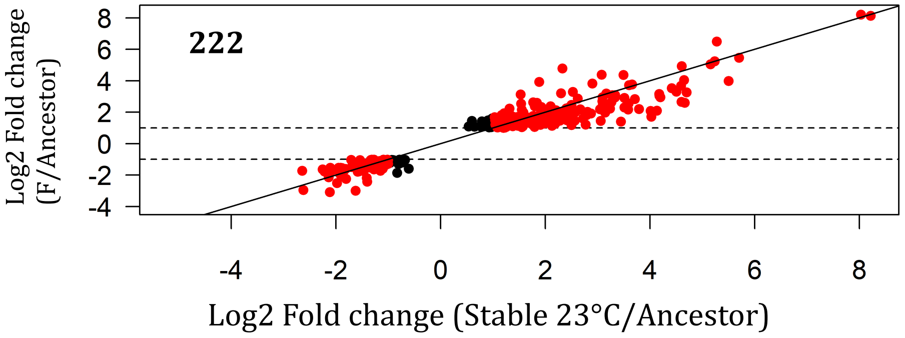
222
Log2 Fold change (F/Ancestor)
Log2 Fold change (Stable 23°C/Ancestor)
8
6
4
2
0
-2
-4
-4
-2
0
2
4
6
8

**H**

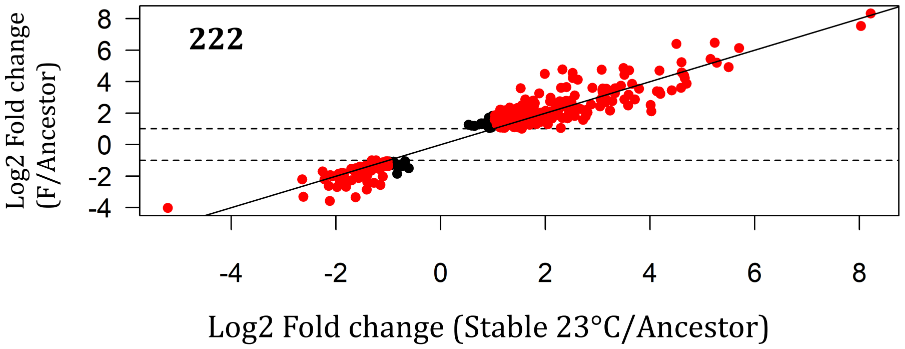
222
Log2 Fold change (F/Ancestor)
Log2 Fold change (Stable 23°C/Ancestor)
8
6
4
2
0
-2
-4
-4
-2
0
2
4
6
8

**Figure S9. Correlation of Log2 fold change between all genes differentially expressed under fluctuation with other evolved lineages. Panels A to D:** genetic background MGGP01; **panels E to H:** genetic background MGGP44 ; results are shown separately for each repeated evolution under fluctuation : see panels **A** and **B**, **C** and **D**, **E** and **F**, **G** and **H**. The number of significant genes in common between the selection regimes is added on each graph. Red points represent genes with a raw Fold change greater than 2.
