## Supplemental Table 6 for "Evolution and plasticity of the transcriptome under temperature fluctuations in the fungal plant pathogen *Zymoseptoria tritici*"

### Additional File 9

**Table S6.** Description of the 63 genes included in the top 8 WGCNA modules. ID: unique gene identifier from the Doe Join Genome Institute, Gene_name: gene number presented in the genome annotation from Grandaubert et al. (2015). Rank: Gene rank within each module depending on their level of connectivity within the module and number of genes in the module; Gene function: function associated to transcription regulation or signal transduction.

| ID | Gene_name | Module | Correlation to the eigengene | Rank | Gene Function |
|---|---|---|---|---|---|
| 43318 | gene_6182 | MGGP01_darkgreen | 0,6857 | 121 / 458 | Transcription |
| 50394 | gene_3427 | MGGP01_darkgreen | 0,7213 | 164 / 458 | Signal transduction |
| 87549 | gene_5500 | MGGP01_darkgreen | 0,8236 | 314 / 458 | Transcription |
| 108064 | gene_7068 | MGGP01_darkgreen | 0,8699 | 388 / 458 | Transcription |
| 83976 | gene_10229 | MGGP01_darkgreen | 0,8952 | 427 / 458 | Transcription |
| 23234 | gene_9490 | MGGP01_red | 0,7501 | 131 / 356 | Transcription |
| 99917 | gene_5845 | MGGP01_red | 0,8435 | 233 / 356 | Signal transduction |
| 47621 | gene_4324 | MGGP01_red | 0,8487 | 240 / 356 | Transcription |
| 86708 | gene_817 | MGGP01_red | 0,8697 | 271 / 356 | Signal transduction |
| 84295 | gene_10819 | MGGP01_red | 0,8846 | 291 / 356 | Signal transduction |
| 110369 | gene_3078 | MGGP01_red | 0,8887 | 298 / 356 | Transcription |
| 60611 | gene_2407 | MGGP01_red | 0,9344 | 344 / 356 | Signal transduction |
| 70309 | gene_9304 | MGGP01_red | 0,9415 | 350 / 356 | Signal transduction |
| 34487 | gene_2657 | MGGP01_red | 0,9710 | 355 / 356 | Signal transduction |
| 66273 | gene_9648 | MGGP01_red | 0,9847 | 356 / 356 | Signal transduction |
| 95583 | gene_2178 | MGGP01_skyblue | 0,7391 | 176 / 456 | Signal transduction |
| 31187 | gene_6733 | MGGP01_skyblue | 0,8530 | 309 / 456 | Transcription |
| 111431 | gene_1815 | MGGP01_skyblue | 0,9361 | 439 / 456 | Signal transduction |
| 93761 | gene_3801 | MGGP44_cyan | 0,6735 | 21 / 224 | Transcription |
| 52472 | gene_6919 | MGGP44_cyan | 0,7294 | 56 / 224 | Signal transduction |
| 92547 | gene_5802 | MGGP44_cyan | 0,8292 | 135 / 224 | Signal transduction |
| 108657 | gene_7540 | MGGP44_cyan | 0,8394 | 143 / 224 | Signal transduction |
| 57142 | gene_9092 | MGGP44_cyan | 0,8620 | 166 / 224 | Transcription |
| 77247 | gene_1282 | MGGP44_cyan | 0,8713 | 175 / 224 | Signal transduction |
| 93065 | gene_2972 | MGGP44_cyan | 0,8787 | 179 / 224 | Signal transduction |

| | | | | | |
|---|---|---|---|---|---|
| 111192 | gene_1442 | MGGP44_cyan | 0,8851 | 187 / 224 | Signal transduction |
| 105024 | gene_27 | MGGP44_grey60 | 0,6285 | 443 / 1509 | Signal transduction |
| 62864 | gene_868 | MGGP44_grey60 | 0,7034 | 672 / 1509 | Signal transduction |
| 65583 | gene_7211 | MGGP44_grey60 | 0,7244 | 767 / 1509 | Transcription |
| 43039 | gene_7058 | MGGP44_grey60 | 0,7707 | 949 / 1509 | Transcription |
| 32659 | gene_9864 | MGGP44_grey60 | 0,7829 | 1004 / 1509 | Transcription |
| 108064 | gene_7068 | MGGP44_grey60 | 0,8208 | 1162 / 1509 | Transcription |
| 74012 | gene_4883 | MGGP44_grey60 | 0,8414 | 1243 / 1509 | Signal transduction |
| 80821 | gene_5349 | MGGP44_grey60 | 0,8506 | 1276 / 1509 | Signal transduction |
| 36765 | gene_5226 | MGGP44_grey60 | 0,8513 | 1278 / 1509 | Signal transduction |
| 77753 | gene_1619 | MGGP44_grey60 | 0,8628 | 1319 / 1509 | Transcription |
| 102849 | gene_10565 | MGGP44_grey60 | 0,8687 | 1342 / 1509 | Transcription |
| 99181 | gene_8278 | MGGP44_grey60 | 0,8713 | 1354 / 1509 | Signal transduction |
| 66328 | gene_9782 | MGGP44_grey60 | 0,8757 | 1372 / 1509 | Signal transduction |
| 88606 | gene_709 | MGGP44_grey60 | 0,8803 | 1380 / 1509 | Signal transduction |
| 79543 | gene_8841 | MGGP44_grey60 | 0,9069 | 1438 / 1509 | Transcription |
| 63560 | gene_2141 | MGGP44_grey60 | 0,9322 | 1480 / 1509 | Transcription |
| 95797 | gene_5348 | MGGP44_grey60 | 0,9481 | 1497 / 1509 | Signal transduction |
| 36575 | gene_4151 | MGGP44_lightcyan | 0,4007 | 95 / 1455 | Signal transduction |
| 106955 | gene_6422 | MGGP44_lightcyan | 0,5008 | 187 / 1455 | Signal transduction |
| 109829 | gene_4859 | MGGP44_lightcyan | 0,6936 | 547 / 1455 | Transcription |
| 89428 | gene_9507 | MGGP44_lightcyan | 0,8009 | 835 / 1455 | Signal transduction |
| 42881 | gene_4920 | MGGP44_lightcyan | 0,8269 | 921 / 1455 | Signal transduction |
| 67073 | gene_10487 | MGGP44_lightcyan | 0,8293 | 932 / 1455 | Transcription |
| 50302 | gene_5238 | MGGP44_lightcyan | 0,8801 | 1146 /1 455 | Signal transduction |
| 101572 | gene_2954 | MGGP44_lightcyan | 0,8844 | 1164 / 1455 | Signal transduction |
| 70921 | gene_4307 | MGGP44_lightcyan | 0,8877 | 1180 / 1455 | Signal transduction |
| 74796 | gene_5093 | MGGP44_lightcyan | 0,8940 | 1210 /1455 | Signal transduction |
| 28576 | gene_6144 | MGGP44_lightcyan | 0,8983 | 1230 / 1455 | Transcription |
| 67977 | gene_3903 | MGGP44_lightcyan | 0,8997 | 1234 / 1455 | Signal transduction |
| 37171 | gene_7192 | MGGP44_lightcyan | 0,9065 | 1265 / 1455 | Transcription |
| 50100 | gene_1575 | MGGP44_lightcyan | 0,9142 | 1295 / 1455 | Signal transduction |
| 65151 | gene_4281 | MGGP44_lightcyan | 0,9304 | 1357 / 1455 | Signal transduction |

| | | | | | |
|---|---|---|---|---|---|
| 46584 | gene_1507 | MGGP44_lightcyan | 0,9350 | 1374 / 1455 | Signal transduction |
| 88104 | gene_5191 | MGGP44_lightcyan | 0,9357 | 1376 / 1455 | Signal transduction |
| 47390 | gene_5057 | MGGP44_lightcyan | 0,9640 | 1435 / 1455 | Transcription |
| 69388 | gene_2933 | MGGP44_lightcyan | 0,9699 | 1442 / 1455 | Transcription |
| 76018 | gene_633 | MGGP44_lightcyan | 0,9701 | 1443 / 1455 | Signal transduction |
