## Supplemental Figures 11 for "Evolution and plasticity of the transcriptome under temperature fluctuations in the fungal plant pathogen *Zymoseptoria tritici*"

**Additional File 10**

**A**

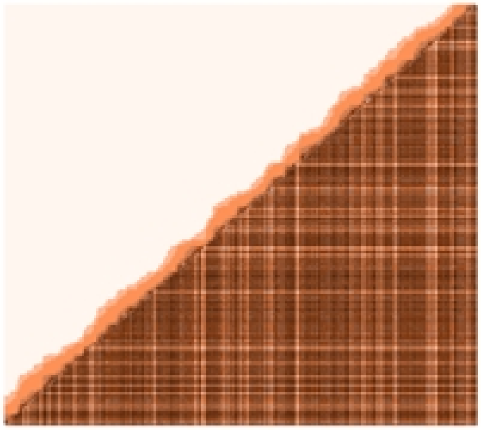

**B**

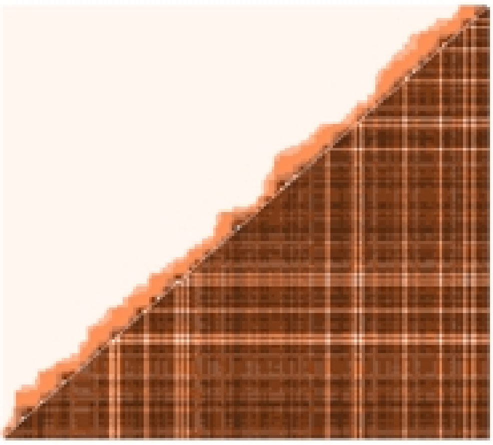

**C**

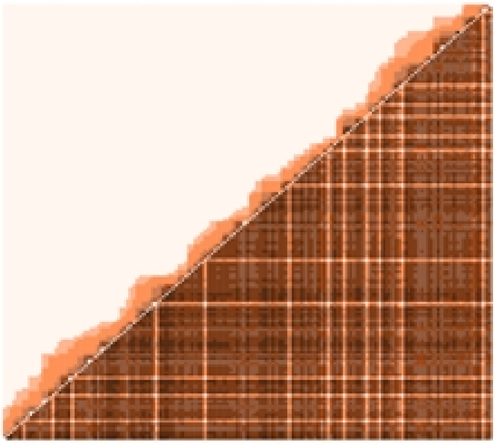

**D**

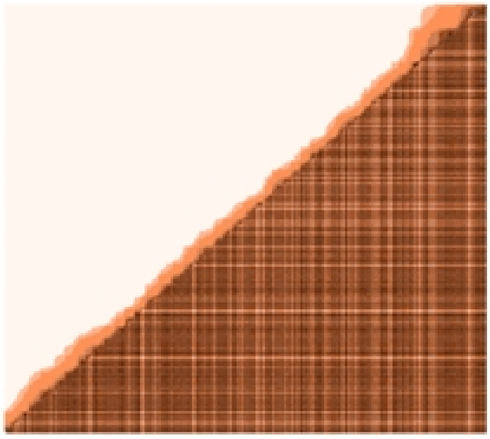

**E**

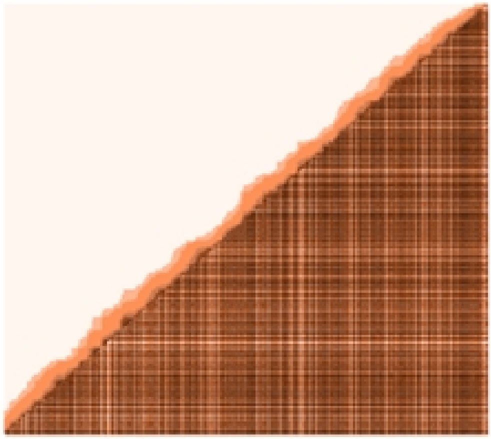

**F**

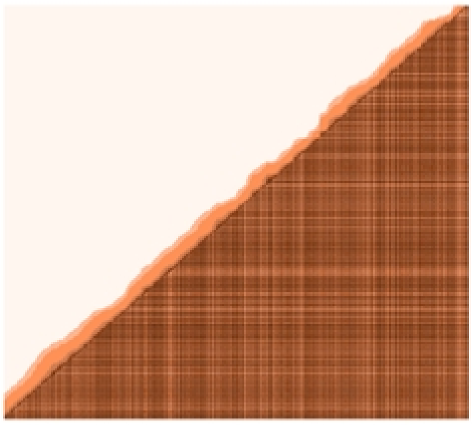

G H

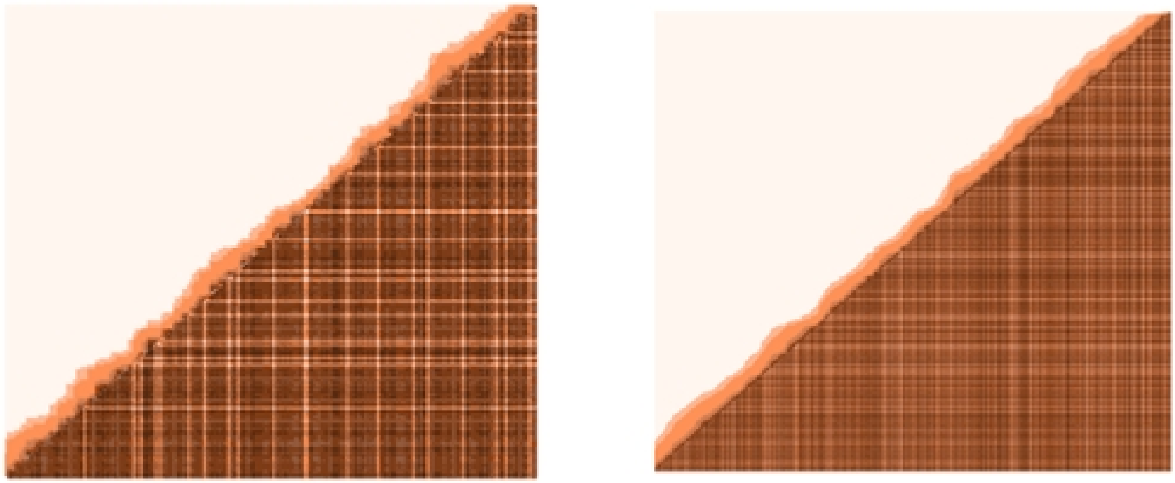

**Figure S11.** Pairwise distance (bp) and pairwise connectivity between genes within each of the top 8 WGCNA modules. Above diagonale: distance between genes; Below diagonale: connectivity. Each graph summarises the relationship among gene for each of the 8 selected modules, named as follow: **A:** red, **B**: purple ; **C:** darkorange ; **D:** darkgreen ; **E:** skyblue, **F:** grey60 ; **G:** cyan ; **H:** lightcyan.
